# Site-Specific Phosphorylation of ZYG-1 Regulates ZYG-1 Stability and Centrosome Number

**DOI:** 10.1101/2023.05.07.539463

**Authors:** Jeffrey C. Medley, Nahyun Yim, Joseph DiPanni, Brandon Sebou, Blake Shaffou, Evan Cramer, Colin Wu, Megan Kabara, Mi Hye Song

**Author notes:** Corresponding author: Mi Hye Song Biological Sciences, Oakland University, 344 Dodge Hall, 118 Library Drive, Rochester, MI 48309, USA.

## Abstract

Spindle bipolarity is critical for genomic integrity. Given that centrosome number often dictates mitotic bipolarity, tight control of centrosome assembly is vital for the fidelity of cell division. The kinase ZYG-1/Plk4 is a master centrosome factor that is integral for controlling centrosome number and is modulated by protein phosphorylation. While autophosphorylation of Plk4 has been extensively studied in other systems, the mechanism of ZYG-1 phosphorylation in *C. elegans* remains largely unexplored. In *C. elegans*, Casein Kinase II (CK2) negatively regulates centrosome duplication by controlling centrosome-associated ZYG-1 levels. In this study, we investigated ZYG-1 as a potential substrate of CK2 and the functional impact of ZYG-1 phosphorylation on centrosome assembly. First, we show that CK2 directly phosphorylates ZYG-1 *in vitro* and physically interacts with ZYG-1 *in vivo.* Intriguingly, depleting CK2 or blocking ZYG-1 phosphorylation at putative CK2 target sites leads to centrosome amplification. In the non-phosphorylatable (NP)-ZYG-1 mutant embryo, the overall levels of ZYG-1 are elevated, leading to an increase in centrosomal ZYG-1 and downstream factors, providing a possible mechanism of the NP-ZYG-1 mutation to drive centrosome amplification. Moreover, inhibiting the 26S proteasome blocks degradation of the phospho-mimetic (PM)-ZYG-1, while the NP-ZYG-1 mutant shows partial resistance to proteasomal degradation. Our findings suggest that site-specific phosphorylation of ZYG-1, partly mediated by CK2, controls ZYG-1 levels via proteasomal degradation, limiting centrosome number. We provide a mechanism linking CK2 kinase activity to centrosome duplication through direct phosphorylation of ZYG-1, which is critical for the integrity of centrosome number.

## Introduction

During cell division, equal distribution of chromosomes into each daughter cell is essential to maintain genomic integrity. As microtubule-organizing centers, centrosomes are critical for establishing mitotic bipolar spindles that ensure accurate transmission of genomic content^1^. Errors in centrosome number disrupt spindle bipolarity, leading to chromosome missegregation, and aberrant centrosomes are linked with human cancers and developmental defects^2,3^. To maintain proper centrosome numbers, centrosomes must duplicate only once per cell cycle through a process tightly regulated and coupled with the cell cycle^1,4^.

Protein phosphorylation has emerged as a critical mechanism that controls the activity and abundance of key centrosome proteins^5^. In humans, the kinase Plk4 is a master regulator of centrosome biogenesis^6^. Plk4 phosphorylates the C-terminal STAN domain of STIL/Ana2/SAS-5, facilitating the direct binding of STIL/Ana2 to SAS-6 and initiating the procentriole formation^7–10^. In a subsequent step, Plk4 phosphorylates another site (S428) of STIL, promoting STIL binding to CPAP/SAS-4^11^. The *C. elegans* ZYG-1, the human Plk4-related kinase, is critical for centrosome duplication^12^. ZYG-1 localizes to the site of procentriole formation^13^ and is necessary for centrosomal recruitment of SAS-5 and SAS-6^14–16^. ZYG-1, independently of its kinase activity, directly binds to SAS-6 to recruit SAS-6 and SAS-5 to nascent centrioles, whereas stable incorporation of SAS-6 into centrioles requires kinase activity of ZYG-1 through an unknown mechanism^17^.

The abundance of centrosome regulators is critical for centrosome number in centrosome assembly. Autophosphorylation and subsequent proteasomal degradation through the SCF-βTrCP/Slimb E3 ubiquitin ligase-dependent pathway have been identified as mechanisms controlling the levels of human Plk4^18,19^. This regulatory mechanism appears to be evolutionarily conserved, involving the SCF-Slimb/βTrCP E3 ubiquitin ligase and the Slimb/βTrCP-binding motif (SBM) in *C. elegans*, *Drosophila*, and human cells^19–22^. In *Drosophila* and human cells, autophosphorylation of the SBM triggers SCF^Slimb/βTrCP^ binding to Plk4, promoting Plk4 degradation^18,23,24^. However, it remains elusive whether autophosphorylation of ZYG-1 *in C. elegans* has a similar role in SCF^Slimb/βTrCP^-mediated degradation of ZYG-1. Several *C. elegans* genes have been identified that modulate ZYG-1 activity in centrosome assembly through multiple regulatory mechanisms, including transcription, RNA binding, proteolysis, and protein phosphorylation^25–29^. Studies in *C. elegans* have shown that protein phosphorylation influences cellular and centrosomal levels of ZYG-1, which is critical for normal centrosome number and microtubule-nucleating activity^25,29–31^. PP2A^SUR-6/B55^ exhibits a strong genetic interaction with ZYG-1 and SAS-5, and PP2A^SUR-6/B55^-dependent dephosphorylation appears to stabilize ZYG-1 and SAS-5 by protecting them from proteasomal destruction^31^. It remains unknown what kinase counteracts PP2A^SUR-6/B55^-mediated dephosphorylation of these centrosome proteins in *C. elegans,* whereas PP2A-dependent dephosphorylation has been proposed to counteract Plk4 autophosphorylation and stabilizes Plk4 in humans and flies^32^.

In *C. elegans,* Casein Kinase II (CK2) has been shown to negatively regulate centrosome duplication^25^. CK2 is a tetrameric holoenzyme comprising two catalytic (CK2α) and two regulatory (CK2β) subunits. The *C. elegans* genome contains the *kin-3* and *kin-10* genes that encode for the CK2α and CK2β subunits, respectively^33,34^, and CK2 functions in various cellular processes^35–43^ including cell division^25^. CK2 is an evolutionarily conserved serine/threonine protein kinase that targets over 500 substrates^44,45^. In human cells, CK2 activity is elevated in both tumor cells and normal proliferating cells^46,47^, and deregulated CK2α activity leads to centrosome amplification, although the precise mechanism is unknown^48^.

In *C. elegans*, CK2 negatively regulates centrosome duplication by controlling centrosome-associated ZYG-1 levels^25^. However, the specific substrate(s) of CK2 and phosphorylation sites involved in centrosome assembly have not yet been identified. In this study, we investigated ZYG-1 as a potential substrate of CK2 and examined the functional impact of site-specific phosphorylation of ZYG-1 on centrosome assembly *in vivo*. Blocking phosphorylation of ZYG-1 at potential CK2 target sites phenocopies loss of CK2 in centrosome assembly. Non-phosphorylatable ZYG-1 is protected from degradation, promoting centrosome amplification. Our findings suggest that multisite phosphorylation of ZYG-1, likely by CK2, regulates ZYG-1 stability and is critical for the proper centrosome number in *C. elegans* embryos.

## Results

### Loss of CK2 Results in Extra Centrosomes

It has been shown that *kin-3(RNAi)*-mediated partial knockdown of CK2 does not significantly impact embryonic development^25,42^. By contrast, embryos further depleted of KIN-3, when animals were treated with *kin-3(RNAi)* for extended hours (40-48 hours), were partially inviable with various cell division phenotypes^25^. Remarkably with further knockdown of KIN-3/CK2, these embryos produced tripolar spindles (**Fig 1A,S1A**). In *C. elegans*, sperm provides a pair of centrioles to the embryo during fertilization. During the first mitosis, the paternal centriole pair segregates and duplicates, then two centrosomes establish bipolar spindles. Live imaging shows that *kin-3(RNAi)-*treated embryos undergo normal cytokinesis at the first mitosis but form tripolar spindles at the second mitosis (**Fig S1A**), suggesting that extra centrosomes likely arise from centrosome overduplication. These observations illustrate that inhibiting CK2 kinase activity leads to centrosome amplification in *C. elegans* embryos, indicating that CK2 kinase activity is critical for the integrity of centrosome number, potentially through phosphorylation of one or more centrosome proteins.

**Figure 1.**
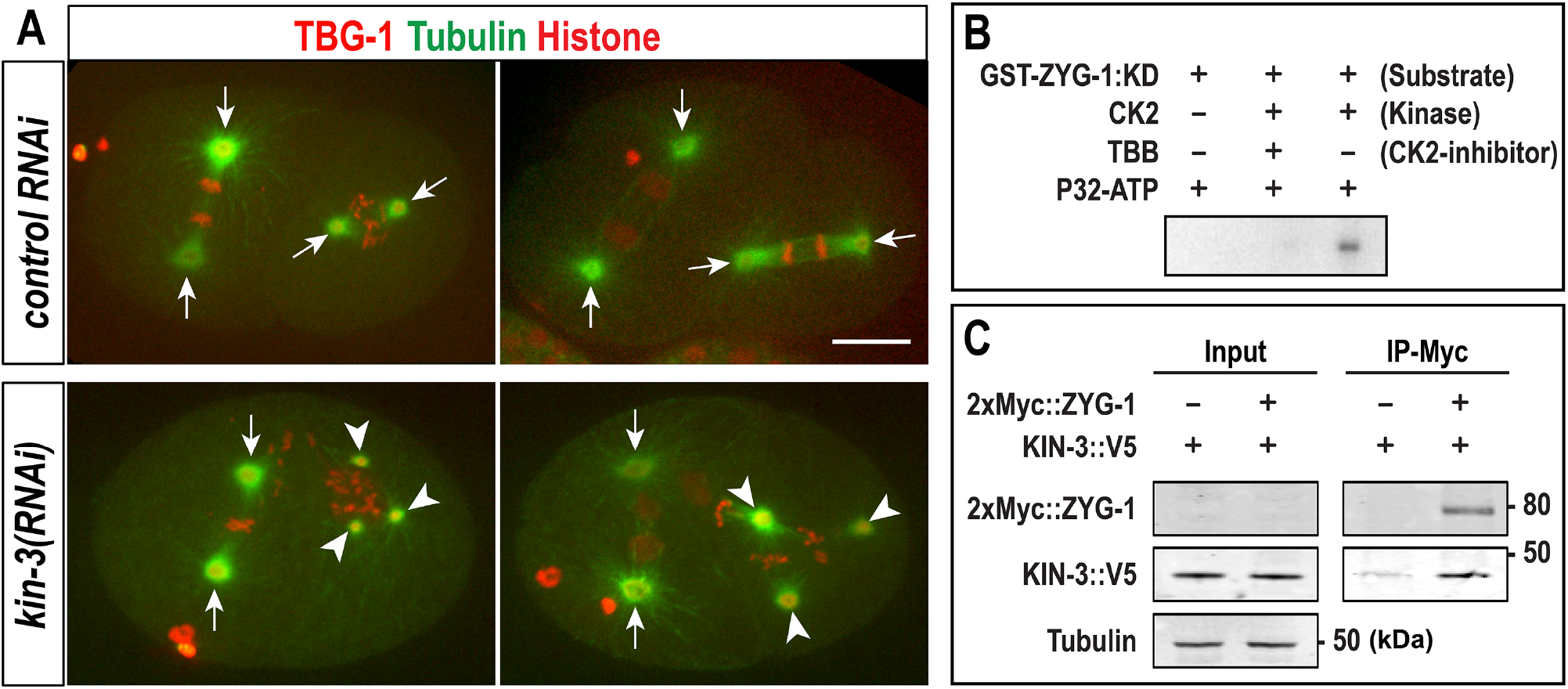
Loss of CK2 leads to extra centrosomes. **(A)** Still images of embryos expressing GFP::β-tubulin, mCherry::TBG-1 (γ-tubulin), and mCherry::histone at the second mitosis. Arrows indicate bipolar spindles, and arrowheads mark tripolar mitotic spindles. Bar, 10µm. **(B)** Autoradiogram of GST-ZYG-1 (kinase-dead: KD) incubated with CK2. **(C)** 2xMyc::ZYG-1 co-precipitates with KIN-3::V5. ∼5% of lysates were loaded in input.

### CK2 phosphorylates ZYG-1 *in vitro*

While CK2 kinase activity influences centrosome assembly in *C. elegans* embryos^25^, we do not know the specific substrate(s) and sites phosphorylated by CK2 critical for centrosome assembly. A compelling candidate is the kinase ZYG-1, a master regulator of centrosome duplication^12^. In *C. elegans*, increased centrosomal ZYG-1 or hyperactive ZYG-1 leads to extra centrosomes in early embryos^20,29^ and seam cells^49^. Because CK2 kinase activity negatively regulates centrosome duplication by controlling ZYG-1 levels at centrosomes^25^, we hypothesized CK2 directly phosphorylates ZYG-1, and that the phosphorylation state of ZYG-1 influences ZYG-1 activity in regulating centrosome assembly. We first tested whether the kinase CK2 can phosphorylate ZYG-1 *in vitro* (**Fig 1B)**. Since the *C. elegans* kinase ZYG-1 is shown to be autophosphorylated *in vitro*^12^, we used the kinase-dead ZYG-1 (KD-ZYG-1^K41M^; ^12^) as a substrate to eliminate auto-phosphorylated ZYG-1 for *in vitro* kinase assay. Autoradiograph illustrates that CK2 phosphorylates KD-ZYG-1^K41M^, and this phosphorylation was blocked by a CK2 kinase inhibitor (TBB). Furthermore, our immunoprecipitation (IP) assays using embryonic lysates suggest a physical interaction between 2xMyc::ZYG-1 and KIN-3::V5 *in vivo*^20,50^ (**Fig 1C**). These results support our hypothesis that CK2 directly phosphorylates ZYG-1.

### CK2 Consensus Sites in the L1 Domain of ZYG-1

CK2 is a serine/threonine protein kinase that favors substrates containing acidic residues downstream of the phosphorylation site^51,52^. Using *in silico* approaches (KinasePhos3.0^53^; and GPS 5.0^54^), we predicted potential phosphorylation sites in ZYG-1, identifying S279, S280, S620, and T621 as having the highest likelihood of CK2 consensus motifs (**Table S1**). These potential CK2 target sites are located in two distinct regions: S279/S280 in the Linker 1 (L1) domain and S620/T621 in the Linker 2 (L2) domain (**Fig 2A**). The sequence alignments illustrate that S279 and S280 exhibit strong evolutionary conservation (**Fig S2A**), but S620 and T621 have weaker conservation (**Fig S2B**) within the *Caenorhabditis* genus. Considering sequence conservation, we focused on S279 and neighboring serine residues within the L1 domain that is critical for ZYG-1 functiosn in centrosome assembly^17,20^. S279 is part of a serine cluster (aa260-285: **Fig 2A**) conforming to the minimal CK2 consensus motif and highly conserved throughout the *Caenorhabditis* genus (**Fig S2A)**. Given that the *C. elegans* ZYG-1 is evolutionarily divergent from Plk4 orthologs, S279 and adjacent serine sites do not appear to be conserved in the human Plk4. However, Plk4 contains several serine/threonine residues that conform to the CK2 consensus motif (**Table S2**). Intriguingly, this ZYG-1 region (aa260-280) coincides with the direct binding domain of ZYG-1 to SAS-6^17^, highlighting the functional significance of this ZYG-1 region in centrosome duplication. Herein, we will refer to this region as ZYG-1:4S.

**Figure 2.**
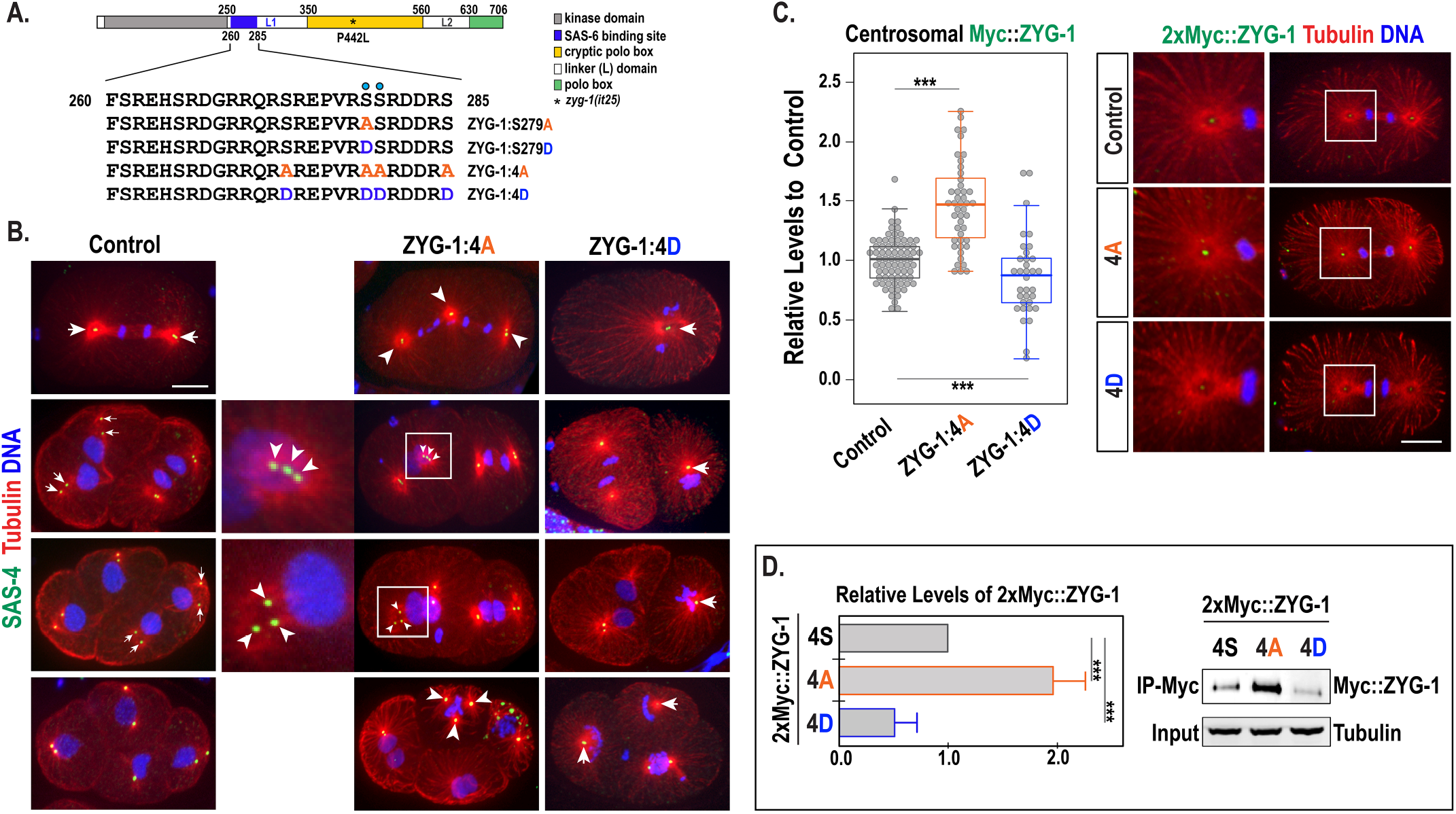
The phosphorylation state of ZYG-1 regulates centrosome number and ZYG-1 stability. **(A)** ZYG-1 protein structure illustrating locations of functional domains, predicted CK2-dependent phosphorylation sites and phospho-mutations. Blue dots indicate S279 and S280. **(B)** Immunostained embryos display abnormal centrosome numbers in ZYG-1 phospho-mutants. Tripolar spindles are observed in ZYG-1:4A mutants and monopolar spindles in ZYG-1:4D mutants. For ZYG-1:4A mutants, insets highlight centrosomes with three centriolar foci stained with SAS-4 at the second telophase. **(C)** (Left) Quantification of centrosomal 2xMyc::ZYG-1 levels during the first anaphase. Each dot represents a centrosome. In the plot, box ranges from the first through third quartile of the data. Thick bar indicates the median. Lines extend to the minimum and maximum data point excluding outliers defined as beyond 1.5 times the interquartile range. (Right) Immunostained embryos. **(D)** (Left) Relative levels of total 2xMyc::ZYG-1 in Myc-pulldown normalized against Tubulin in input (mean±s.d.). (Right) Myc-IP of 2xMyc::ZYG-1 embryonic lysates. ∼5% of total lysates were loaded in input. (**B,C**) Insets magnified 4-fold. Bar, 10µm. **(C,D)** \*\*\**p*<0.001 (two-tailed unpaired *t*-tests).

To test whether CK2 phosphorylates ZYG-1 at these sites, we performed *in vitro* kinase assay using the ZYG-1 peptides (aa260-285) containing four serine residues as a substrate (**Fig S2C**). Autoradiogram of thin layer chromatography (TLC) revealed that CK2 phosphorylates the wild-type ZYG-1 peptide but not the ZYG-1:4A peptide where four sites (S273, S279, S280, and S285) were replaced with alanine. By contrast, the CK2 inhibitor TBB reduced CK2-dependent phosphorylation of the ZYG-1 peptide. Mass spectrometry analysis of *in vitro* kinase reactions using the ZYG-1 peptide substrate and CK2 confirmed the reproducible phosphorylation of at least one serine residue (S279 or S280) within this ZYG-1 peptide (**Fig S2D**^55^). Our data suggest CK2 directly phosphorylates at least one serine residue in the ZYG-1 L1 domain.

### Non-phosphorylatable ZYG-1 Mutation Leads to Centrosome Amplification

To address the functional impacts of site-specific phosphorylation of ZYG-1 *in vivo*, we used CRISPR/Cas9 editing to mutate S279 to a non-phosphorylatable (NP) alanine, termed ZYG-1^S279A^ and a phospho-mimetic (PM) aspartate, termed ZYG-1^S279D^ at the endogenous locus (**Fig 2A**). In parallel, we simultaneously mutated four serine residues (S273, S279, S280 and S285) to alanine, termed ZYG-1^4A^, or aspartate, termed ZYG-1^4D^. The homozygous ZYG-1 phospho-mutants we generated were mostly viable, with ZYG-1^4A^ and ZYG-1^4D^ mutants producing a low rate of embryonic lethality (**Table 1**). Since *kin-3(RNAi)*-mediated inhibition of CK2 leads to tripolar spindle formation (**Fig 1A**), we speculated that mutating a subset of CK2 target sites to non-phosphorylatable alanine (NP-ZYG-1 mutations) should partially phenocopy loss of CK2, including extra centrosomes. We first observed centrosome behavior to determine how the phosphorylation state of ZYG-1 affected centrosome duplication (**Fig 2B,S3**). Intriguingly, we observed extra centrosomes in ZYG-1^4A^ mutant embryos, starting at the first mitosis and later cell divisions (∼30%, n>100 embryos). Extra centrosomes resemble phenotypes arising from Plk4 overexpression^6^, implying that the ZYG-1^4A^ mutation upregulates ZYG-1 activity and hyperactive ZYG-1 leads to centrosome amplification. Furthermore, we observed various cell division phenotypes in ZYG-1^4A^ mutant embryos (**Fig S3A**), including DNA missegregation, cytokinesis failure, detached centrosomes, and expanded PCM morphology, similar to phenotypes observed in embryos depleted of CK2^25^. These results show that the NP-ZYG-1^4A^ mutation phenocopies loss of CK2 during *C. elegans* embryogenesis. In contrast, the ZYG-1^4D^ mutation caused monopolar spindle formation in one-cell and later-stage embryos (∼30%, n>100) (**Fig 2B,S3C**), indicating that the PM-ZYG-1^4D^ mutation downregulates ZYG-1 activity and impairs centrosome duplication during spermatogenesis and mitotic divisions.

**Table 1.** Genetic Analysis.

| Strain | °C | % Embryonic Viability<br>(ave ± s.d.) | n<br>(progeny) |
| --- | --- | --- | --- |
| wild-type (N2) | 22.5 | 100 ± 0 | 1803 |
| ZYG-1<sup>S279A</sup> | | 98.7 ± 2.5 | 1542 |
| ZYG-1<sup>4A</sup> | | 90.4 ± 5.0 | 2052 |
| ZYG-1<sup>S279D</sup> | | 99.0 ± 3.3 | 1669 |
| ZYG-1<sup>4D</sup> | | 98.6 ± 1.7 | 1718 |
| *zyg-1(it25)* | | 8.8 ± 8.2 | 1275 |
| ZYG-1<sup>S279A</sup>: *zyg-1(it25)* | | 31.1 ± 19.9 | 1301 |
| ZYG-1<sup>4A</sup>: *zyg-1(it25)* | | 96.1 ± 3.6 | 3426 |
| ZYG-1<sup>S279D</sup>: *zyg-1(it25)* | | 2.5 ± 2.8 | 1401 |
| ZYG-1<sup>4D</sup>: *zyg-1(it25)* | | 1.9 ± 3.4 | 1529 |
| wild-type (N2) | 23.0 | 100 ± 0 | 400 |
| ZYG-1<sup>S279A</sup> | | 100 ± 0 | 504 |
| ZYG-1<sup>4A</sup> | | 87.7 ± 6.4 | 457 |
| ZYG-1<sup>S279D</sup> | | 97.6 ± 1.2 | 573 |
| ZYG-1<sup>4D</sup> | | 99.1 ± 0.6 | 336 |
| *zyg-1(it25)* | | 0.3 ± 0.7 | 716 |
| ZYG-1<sup>S279A</sup>: *zyg-1(it25)* | | 3.7 ± 3.1 | 301 |
| ZYG-1<sup>4A</sup>: *zyg-1(it25)* | | 19.7 ± 6.2 | 514 |
| ZYG-1<sup>S279D</sup>: *zyg-1(it25)* | | 0 ± 0 | 239 |
| ZYG-1<sup>4D</sup>: *zyg-1(it25)* | | 0 ± 0 | 494 |

### Non-phosphorylatable ZYG-1 Leads to Elevated ZYG-1 Levels

Since inhibiting CK2 kinase activity leads to elevated centrosomal ZYG-1^25^, we examined how the NP-and PM-ZYG-1 mutations affected ZYG-1 levels at centrosomes. To facilitate this assay, we introduced an N-terminal 2xMyc tag at the endogenous *zyg-1* locus by CRISPR/Cas9 editing^20^. By immunostaining embryos with anti-Myc, we quantified the fluorescence intensity of centrosome-associated 2xMyc::ZYG-1 during the first mitosis (**Fig 2C**). The NP-ZYG-1^4A^ still localizes to centrosomes, indicating alanine replacements at these serine sites did not impair ZYG-1 loading to centrosomes. Quantitative immunofluorescence (IF) revealed that ZYG-1^4A^ mutant embryos exhibit significantly increased 2xMyc::ZYG-1 signals at centrosomes (1.47±0.45 fold, *p*<0.001), while ZYG-1^4D^ mutants showed decreased levels (0.81±0.28 fold, *p*<0.01), compared to wild-type controls (1.00±0.20 fold). Similar trends were observed at the first metaphase (**Fig S4A**). Our data illustrate that the phosphorylation state of ZYG-1 affects its protein levels at centrosomes.

Increased levels of cellular ZYG-1 may account for elevated centrosomal ZYG-1 in NP-ZYG-1 mutants. Following Myc-pulldown using embryonic lysates expressing 2xMyc::ZYG-1, we could detect Myc-tagged ZYG-1, with significantly increased levels (1.98±0.51 fold, *p<*0.0001, n=6) of 2xMyc::ZYG-1 in ZYG-1^4A^ embryos, and reduced levels (0.51±0.19 fold, *p<*0.0001, n=6) in ZYG-1^4D^ mutants (**Fig 2D**). Our quantitative immunoblot supports a model where NP-ZYG-1 is hyperstabilized, leading to increased levels of cellular ZYG-1, thereby more ZYG-1 loaded to centrosomes.

### Non-phosphorylatable ZYG-1 Restores Centrosome Duplication in *zyg-1(it25)*

Depleting CK2 restores centrosome duplication and embryonic viability to *zyg-1(it25)* mutants^25^. We thus asked if the NP-ZYG-1 mutation, mimicking loss of CK2, could produce similar effects on *zyg-1(it25)* mutants. In this scenario, the NP-ZYG-1 mutation should restore embryonic viability and centrosome duplication to *zyg-1(it25)* mutants. To test this, we used CRISPR/Cas9 editing and introduced equivalent phospho-mutations to *zyg-1(it25)* mutants at the endogenous locus. In the temperature-sensitive *zyg-1(it25)* mutant embryos grown at the restrictive temperature 24°C, centrosome duplication fails during the first cell cycle, resulting in monopolar spindles at the second mitosis and 100% embryonic lethality^12^. Remarkably, the ZYG-1^4A^ mutation led to a significant restoration (>10-fold) of embryonic viability, and the ZYG-1^S279A^ mutation produced a relatively moderate but significant restoration (3.5-fold) to *zyg-1(it25)* mutants at semi-restrictive temperature conditions (**Table 1, Fig 3A**). By contrast, the ZYG-1^4D^ and ZYG-1^S279D^ mutations produced opposite effects on *zyg-1(it25)* mutants (**Table 1, Fig 3A**). These results suggest that the NP-ZYG-1 mutation upregulates ZYG-1 activity, compensating for the hypomorphic ZYG-1 function in *zyg-1(it25)* mutants, while the PM-ZYG-1 mutations have the opposite effect.

**Figure 3.**
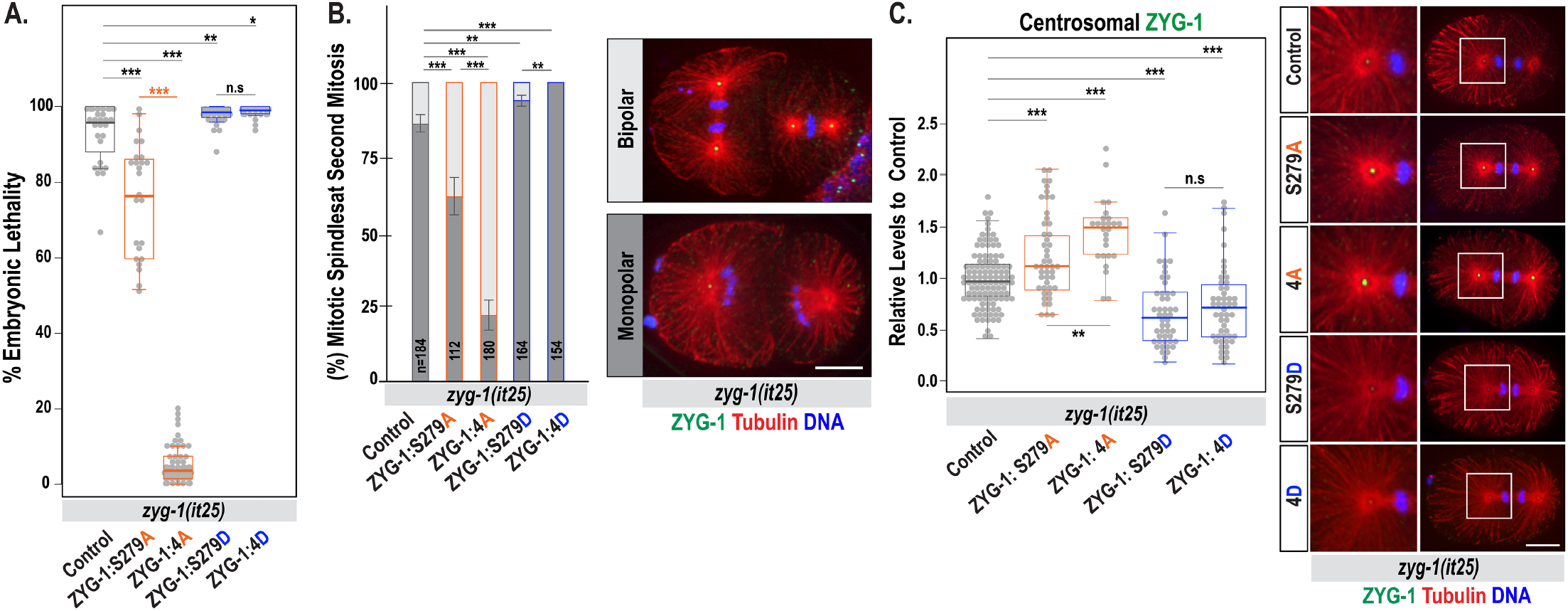
Multisite phosphorylation of ZYG-1 regulates ZYG-1 activity and centrosome duplication. **(A)** Embryonic viability at 23°C (Table 1). Each dot represents a hermaphrodite. **(B)** (Left) Quantification of mitotic spindles at the second mitosis at 23°C (mean±s.d.). n is the number of blastomeres. (Right) Immunostained embryos undergoing second mitosis illustrate monopolar (dark grey) or bipolar (light grey) spindles. **(C)** (Left) Quantification of centrosomal ZYG-1 levels at the first anaphase. Each dot represents a centrosome. (Right) Embryos stained for ZYG-1 at the first anaphase. Insets magnified 4-fold. **(B,C)** Bar, 10µm. **(A-C)** n.s *p*>0.05, \**p*<0.05, \*\**p*<0.01, \*\*\**p*<0.001 (two-tailed unpaired *t*-tests).

Given that ZYG-1 is essential for centrosome duplication, restoring embryonic viability to *zyg-1(it25)* mutants likely results from successful centrosome duplication. To examine how ZYG-1 phospho-mutations affected centrosome duplication in the *zyg-1(it25)* mutant, we scored for bipolar spindle formation during the second mitosis (**Fig 3B**). Consistent with the genetic suppression, both NP-ZYG-1^S279A^ and ZYG-1^4A^ mutations restored bipolar spindles to *zyg-1(it25)* mutants (13.8±6.2%), with the ZYG-1^4A^ mutation (77.7±10%) exhibiting a higher rate of bipolar spindles than the ZYG-1^S279A^ mutation (38.1±12%). Conversely, both PM-ZYG-1 mutations (ZYG-1^S279D^ and ZYG-1^4D^) decreased bipolar spindle formation (6±3.6% and 0%, respectively). Similar to loss of CK2, the NP-ZYG-1 mutations appear to enhance ZYG-1 activity, thereby restoring centrosome duplication and embryonic viability to *zyg-1(it25)* mutants, while the PM-ZYG-1 mutations diminish ZYG-1 activity and aggravate *zyg-1(it25)* phenotypes.

Increased levels of centrosomal ZYG-1 in ZYG-1^4A^ mutants may explain suppression of *zyg-1(it25)* phenotypes by the NP-ZYG-1 mutation. To directly test this, we compared centrosomal ZYG-1 levels in *zyg-1(it25)* mutant backgrounds (**Fig 3C**). Quantitative IF using anti-ZYG-1^56^ illustrate significantly increased levels of centrosomal ZYG-1 in ZYG-1^4A^: *zyg-1(it25)* (1.45±0.33 fold, *p*<0.001) and ZYG-1^S279A^: *zyg-1(it25)* (1.26±0.43 fold, *p*<0.001) embryos, compared to *zyg-1(it25)* controls (1.00±0.26 fold) during the first anaphase. Notably, the ZYG-1^4A^ mutation produced a stronger impact on centrosomal ZYG-1 levels than the ZYG-1^S279A^ mutation, consistent with more robust *zyg-1* suppression by ZYG-1^4A^ than the ZYG-1^S279A^ mutation (**Fig 3A,B and Table 1**). Conversely, reduced centrosomal ZYG-1 levels were observed in the ZYG-1^4D^: *zyg-1(it25)* (0.75±0.42-fold, *p*<0.001) and ZYG-1^S279D^: *zyg-1(it25)* (0.67±0.33-fold, *p*<0.001) mutants at the first anaphase. Together, our results suggest that phosphorylation of ZYG-1 at multiple sites regulates centrosomal ZYG-1 levels, thereby influencing ZYG-1 activity in centrosome assembly.

### ZYG-1 Phosphorylation Affects Centrosomal Loading of Downstream Centrosome Factors

In the molecular hierarchy of centrosome assembly, SPD-2 acts upstream and promotes centrosomal targeting of ZYG-1 via direct electrostatic interaction^57^, and ZYG-1 is required for centrosomal recruitment of downstream centrosome factors, SAS-6, SAS-5 and SAS-4^14,17^. Then, elevated centrosomal ZYG-1 in ZYG-1^4A^ mutants may enhance centrosomal recruitment of downstream factors, leading to centrosome amplification. Alternatively, the phosphorylation state of ZYG-1 could affect its interaction with other centrosome factors positively or negatively.

Quantitative IF during the first anaphase reveals that NP-ZYG-1^4A^ mutant embryos exhibit increased levels of centrosomal SAS-6 (1.26±0.31 fold, *p*<0.001), whereas ZYG-1^4D^ mutant embryos show reduced SAS-6 levels (0.73±0.17 fold, *p*<0.01), compared to wild-type controls (1.00±0.28 fold) (**Fig 4A**). Increased SAS-6 levels in NP-ZYG-1^4A^ mutant centrosomes might result from elevated centrosomal ZYG-1. Alternatively, the phosphorylation state of ZYG-1 could affect the ZYG-1-SAS-6 binding affinity through charge changes, as the phosphorylation cluster (ZYG-1:4S) coincides with the ZYG-1 domain that directly interacts with SAS-6 through electrostatic attraction^17^. To investigate this, we performed quantitative immunoblot analysis following Myc-IP to compare the relative levels of SAS-6 associated with 2xMyc::ZYG-1 (**Fig S2E**). Both Myc-IPs of ZYG-1^4A^ and ZYG-1^4D^ mutants show a higher ratio of SAS-6 to 2xMyc::ZYG-1 than controls, implying that the phosphorylation states of ZYG-1 do not adversely affect the ZYG-1-SAS-6 binding affinity. Thus, elevated centrosomal ZYG-1 in ZYG-1^4A^ mutants seems more likely to further enhance SAS-6 recruitment to centrosomes.

**Figure 4.**
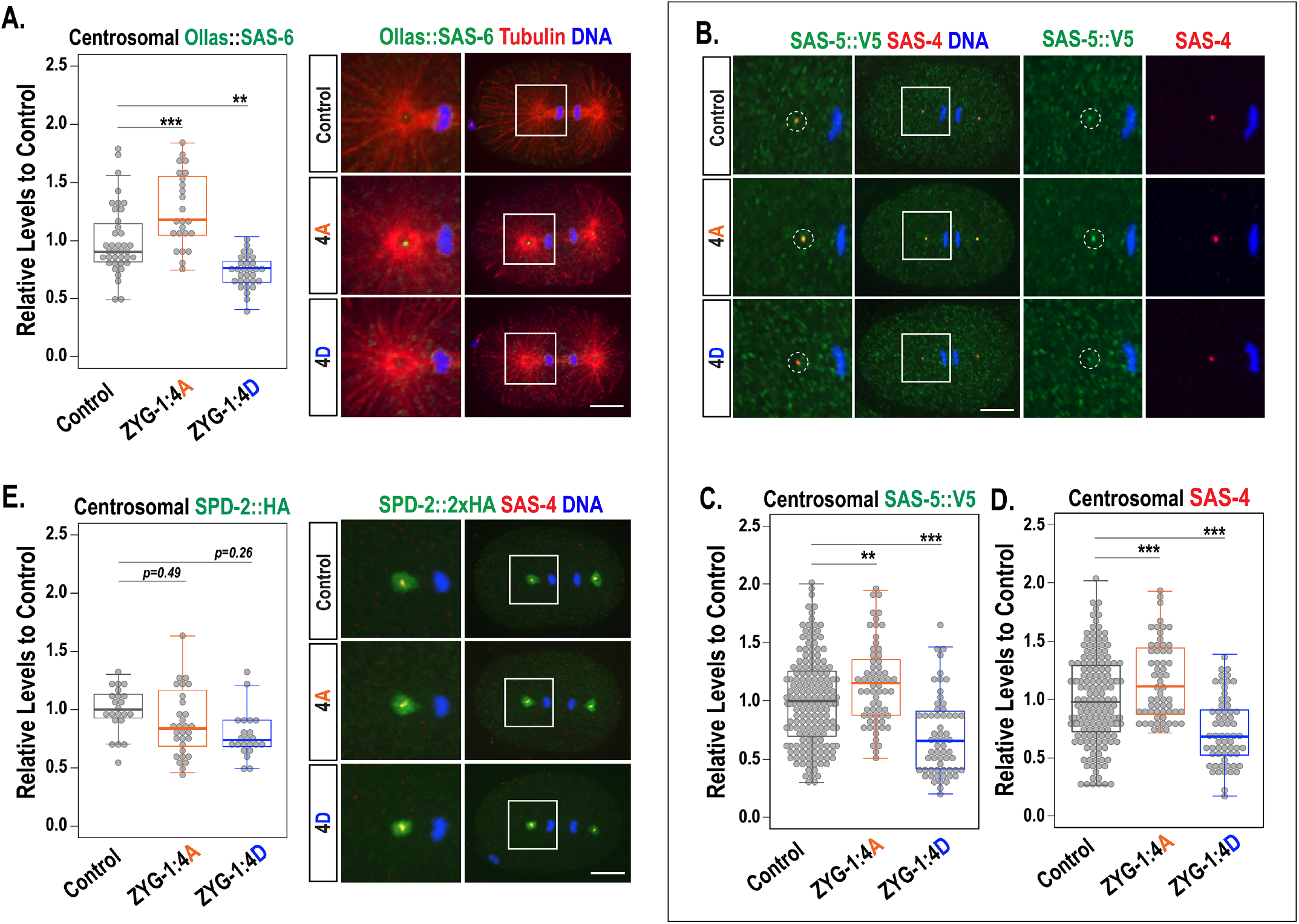
The phosphorylation state of ZYG-1 influences centrosomal levels of downstream factors. ZYG-1 phospho-mutant embryos stained for **(A)** Ollas::SAS-6, **(B)** SAS-5::V5 and SAS-4, **(E)** SPD-2::2xHA. **(B)** The dotted circles highlight co-localization of SAS-5::V5 and SAS-4. **(A,B,E)** Insets magnified 4-fold. Bar, 10µm. **(A,C-E)** Plots show quantification of centrosomal proteins at the first mitotic anaphase. \*\**p*<0.01, \*\*\**p*<0.001 (two-tailed unpaired *t*-tests).

ZYG-1 is also required for centrosomal SAS-5 loading, and SAS-5 and SAS-6 are co-dependent for centrosomal localization^15^. Consistent with SAS-6, we observed significantly increased SAS-5 in ZYG-1^4A^ mutant centrosomes (1.20±0.43 fold, *p*<0.001), but decreased levels in ZYG-1^4D^ centrosomes (0.73±0.29 fold, *p*<0.001), compared to controls (1.00 ± 0.40 fold) (**Fig 4B,C**). Similar trends were observed for centrosomal SAS-4, with a significant increase in ZYG-1^4A^ mutants (1.20±0.43 fold, *p*<0.001) and a decrease in ZYG-1^4D^ mutants (0.73±0.29 fold, *p*<0.001) (**Fig 4B,D**). We also quantified centrosomal levels of SAS-6, 5, and 4 at the first metaphase, showing similar trends in ZYG-1 phospho-mutants (**Fig S4B-D**).

We next asked how ZYG-1 phosphorylation influences centrosomal levels of SPD-2 that functions upstream of ZYG-1 and is required for centrosomal recruitment of ZYG-1^14,58^. The interaction between SPD-2 and ZYG-1 relies on electrostatic interactions between the N-terminal SPD-2 acidic region and the ZYG-1 cryptic polo box (CPB) basic patch (aa350-560^57^). Since the ZYG-1 L1 domain bridges the kinase and CPB domains, phosphorylation of ZYG-1:4S may affect the interaction between ZYG-1 and SPD-2, influencing centrosomal recruitment of ZYG-1. To assess this, we quantified the total centrosomal signals of SPD-2::2xHA in ZYG-1 phospho-mutants using strains expressing SPD-2::2xHA at the endogenous locus. In contrast to ZYG-1 downstream factors, we found no significant changes in centrosomal SPD-2 levels in ZYG-1^4A^ (1.11±0.58 fold, *p*=0.49) and ZYG-1^4D^ mutants (1.08±0.56 fold, *p*=0.26) compared to controls (1.00 ± 0.19 fold) at the first anaphase (**Fig 4E**), and similar trends at the metaphase (**Fig S4E,F**). Furthermore, the phosphorylation state of ZYG-1 did not significantly affect centrosomal levels of the PCM component TBG-1/γ-tubulin during the first mitosis (**Fig S4G**). Given ZYG-1 acts downstream of SPD-2 in centrosome assembly, our results are consistent with a genetic hierarchy where ZYG-1 phospho-mutations had no significant effect on centrosomal SPD-2.

Collectively, our data illustrate that the phosphorylation state of ZYG-1 affects centrosomal recruitment of downstream centrosome proteins, partially explaining how phosphorylation of ZYG-1 influences centrosome number in the *C. elegans* embryo.

### CK2-dependent Phosphorylation of ZYG-1

If CK2 targets one or more serine residues in the ZYG-1:4S region (**Fig 2A**), ZYG-1 phospho-mutants should be less responsive to CK2 depletion than controls. Conversely, if CK2 targets none of the serine residues in this region, ZYG-1 phospho-mutants should be as responsive to CK2 depletion as controls. For the latter, combining CK2 depletion with the ZYG-1^4A^ mutation should show an additive effect. To test this, we used *kin-3(RNAi)* to knockdown KIN-3, the catalytic subunit of CK2, and assayed for bipolar spindle formation in *zyg-1(it25)* genetic backgrounds at the second mitosis (**Fig 5A,B**). As shown in *zyg-1(it25)* controls^25^, *kin-3(RNAi)* significantly increased bipolar spindle formation by 6.7-fold, compared to control RNAi. For ZYG-1^4A^: *zyg-1(it25)* mutants, control RNAi and *kin-3(RNAi)*-treated embryos produced 75±7.8% and 98±2.4% bipolarity, respectively, showing that *kin-3(RNAi)* only caused a slight 1.3-fold increase in bipolarity compared to control RNAi. In ZYG-1^4D^: *zyg-1(it25)* mutants, control RNAi produced nearly zero bipolarity (0.4±1.1%), while *kin-3(RNAi)* led to 16.1±3.3% bipolarity, much lower than expected to be additive (∼44%). These results indicate that the PM-ZYG-1^4D^ mutant is partially resistant to CK2 knockdown, further supporting that CK2 phosphorylates one or more serine residues in the ZYG-1:4S region.

**Figure 5.**
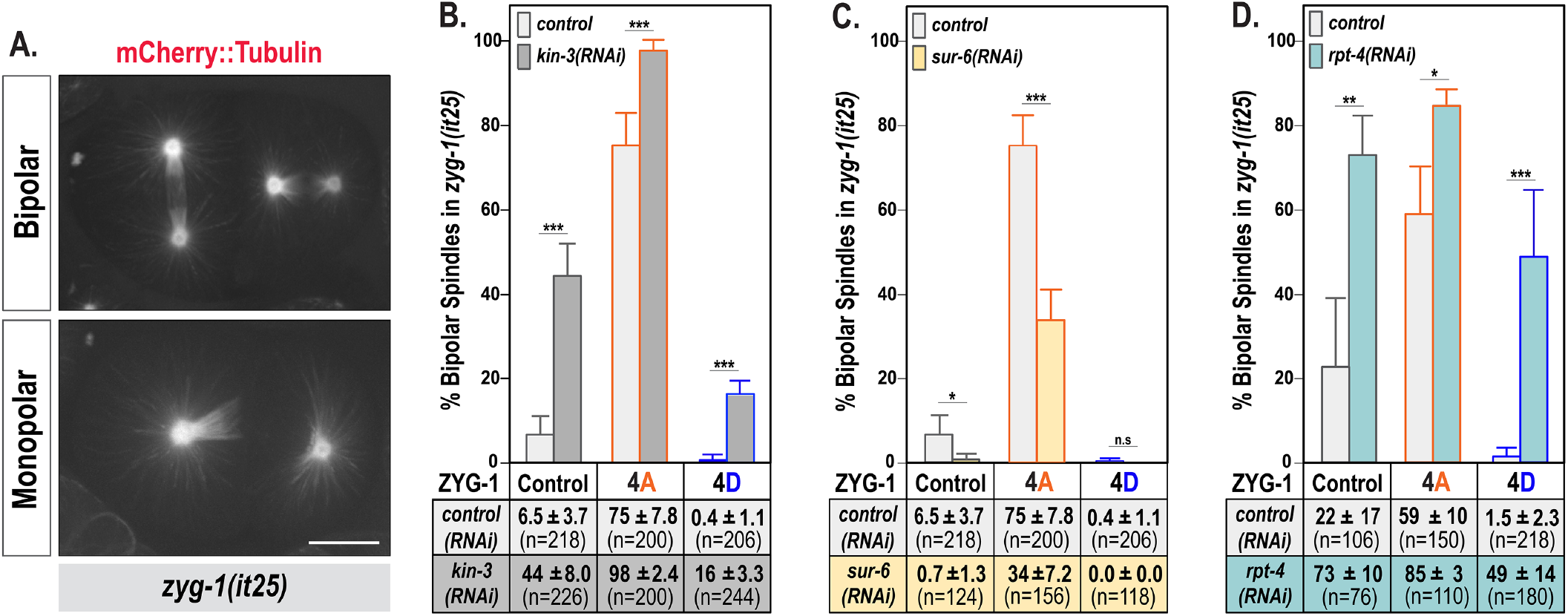
Phosphorylation of ZYG-1 promotes proteasomal degradation of ZYG-1. **(A)** *zyg-1(it25)* mutant embryos expressing mCherry::Tubulin undergoing second mitosis illustrate bipolar and monopolar spindles. Bar, 10µm. **(B-D)** Quantification of bipolar spindle formation at the second mitosis in *zyg-1(it25)* phospho-mutants at 24°C after knocking down **(B)** KIN-3, **(C)** SUR-6, or **(D)** RPT-4. (mean± s.d.). n is the number of blastomeres. \**p*<0.05, \*\**p*<0.01, \*\*\**p*<0.001 (two-tailed unpaired *t*-tests).

Additionally, if the ZYG-1:4S region represents the only target sites for CK2 phosphorylation, *kin-3(RNAi)* should not affect % bipolarity in ZYG-1^4A^: *zyg-1(it25)* mutants. However, *kin-3(RNAi)* led to a moderate but noticeable increase of bipolarity in both *zyg-1(it25)* phospho-mutants (**Fig 5B**), implying that CK2 likely targets additional ZYG-1 residues or other substrates that influence centrosome assembly directly or indirectly.

### PP2A^SUR-6/B55^ May Counteract CK2 to Balance the Phosphorylation State of ZYG-1

In *C. elegans*, the kinase CK2 and the phosphatase PP2A^SUR-6/B55^ play opposing roles in centrosome duplication; CK2 as a negative regulator^25^ and PP2A^SUR-6/B55^ as a positive regulator^30,31^. Inhibiting CK2 increases centrosomal ZYG-1 levels^25^, whereas loss of PP2A^SUR-6/B55^ decreases cellular ZYG-1 levels^31^, suggesting that CK2 and PP2A^SUR-6/B55^ counteract to regulate centrosome assembly by controlling ZYG-1 levels. Intriguingly, serine residues in the ZYG-1:4S conform to both CK2 phosphorylation sites^52^ and minimal PP2A^SUR-6/B55^ consensus motifs^59,60^.

Loss of PP2A^SUR-6/B55^ has been shown to enhance monopolar spindles in *zyg-1(it25)* mutants, in contrast to the impact of CK2 depletion^25,31^. A recent study in *C. elegans* also showed that *kin-3* suppresses *sur-6,* encoding the SUR-6/B55 regulatory subunit of PP2A, in centrosome duplication^61^. Our results indicate that the PM-ZYG-1^4D^ mutation enhances monopolar spindles in *zyg-1(it25)* mutants, whereas the NP-ZYG-1^4A^ mutation has the opposite effect (**Fig 3B**), which led us to speculate that PP2A^SUR-6/B55^ might dephosphorylate the ZYG-1:4S and counteract CK2 to balance the phosphorylation state of ZYG-1. If PP2A^SUR-6/B55^ dephosphorylates serine residues in the ZYG-1:4S region, the phospho-mutants should exhibit partial resistance to SUR-6 inhibition. We treated animals with *sur-6(RNAi)* to partially deplete PP2A^SUR-6/B55^ and observed centrosome duplication (**Fig 5D**). Consistent with previous findings^31^, *sur-6(RNAi)* decreased bipolarity by 9-fold in the *zyg-1(it25)* control, compared to control RNAi. In the ZYG-1^4A^: *zyg-1(it25)* mutants, *sur-6(RNAi)* produced only a minor decrease in bipolarity (34±7.2%, 2-fold) relative to control RNAi (75±7.8%), indicating partial resistance of the ZYG-1^4A^ to PP2A^SUR-6/B55^ knockdown. Thus, PP2A^SUR-6/B55^ likely targets the ZYG-1:4S region. Our collective data suggest that CK2 and PP2A^SUR-6/B55^ target one or more shared serine residues in the ZYG-1:4S region and counteract to balance the phosphorylation state of ZYG-1, providing a regulatory mechanism to ensure that ZYG-1 levels remain above the threshold to support centrosome duplication.

### Phosphorylation of ZYG-1 Promotes Proteasomal Degradation

Our data support a model where the phosphorylation state of ZYG-1 regulates overall ZYG-1 levels, likely by influencing protein stability. One possible mechanism is that phosphorylated ZYG-1 in the ZYG-1:4S undergoes proteasomal degradation, while dephosphorylated ZYG-1 is protected from degradation and stabilized. To test whether ZYG-1 phosphorylation in the ZYG-1:4S promotes proteasomal degradation, we treated ZYG-1 phospho-mutant *zyg-1(it25)* animals with *rpt-4(RNAi)* to partially deplete proteasome activity under conditions that allow completion of meiotic and early mitotic cycles^31^. We then examined how inhibiting the 26S proteasome affected the activity of ZYG-1^4A^ and ZYG-1^4D^ in centrosome duplication (**Fig 5A,D**). If *rpt-4(RNAi)* partially blocked degradation of the phospho-mimetic ZYG-1 (ZYG-1^4D^), *rpt-4(RNAi)* should protect the unstable ZYG-1^4D^ partially from degradation, and then stabilized ZYG-1^4D^ should restore bipolarity in ZYG-1^4D^: *zyg-1(it25)* embryos.

First, we confirmed that in *zyg-1(it25)* controls, *rpt-4(RNAi)* significantly increased bipolar spindle formation by 3.4-fold, compared to control RNAi, consistent with previous work^20^. As expected, in ZYG-1^4D^: *zyg-1(it25)* mutants, compared to control RNAi, *rpt-4(RNAi)* produced a stark increase in bipolar spindles (49±14%, 33.5 fold), close to the percentage of bipolarity observed in the ZYG-1^4A^: *zyg-1(it25)* mutants (59±10%). By contrast, in ZYG-1^4A^: *zyg-1(it25)* mutants, *rpt-4(RNAi)* produced only a minor increase (1.4-fold) compared to control RNAi. Together, *rpt-4(RNAi)* leads to a remarkable stabilization of the ZYG-1^4D^ but a minor effect on the ZYG-1^4A^, suggesting that inhibiting the 26S proteasome blocks proteasomal degradation of ZYG-1^4D^, whereas ZYG-1^4A^ is partially resistant to proteasomal degradation. It seems plausible that phosphorylated ZYG-1 is targeted for proteasomal destruction, triggered by CK2-dependent phosphorylation at ZYG-1:4S. These results support a model where site-specific phosphorylation of ZYG-1 by CK2 regulates ZYG-1 stability through the 26S proteasome, providing a critical mechanism for maintaining the integrity of centrosome number in *C. elegans* embryos.

## Discussion

### CK2 kinase activity is critical for the integrity of centrosome number

In this study, we demonstrate that inhibiting CK2 leads to centrosome amplification (**Fig 1A**), highlighting the critical role of CK2 kinase activity in maintaining the proper centrosome number. Since CK2 negatively regulates centrosome duplication by controlling ZYG-1 levels at centrosomes^25^, we hypothesized that CK2 directly phosphorylates ZYG-1 and ZYG-1 phosphorylation by CK2 influences ZYG-1 activity in centrosome assembly. First, we show that CK2 directly phosphorylates kinase-dead ZYG-1 *in vitro,* and physically interacts with ZYG-1 *in vivo* (**Fig 1B,C**). To investigate the functional consequences of ZYG-1 phosphorylation at the putative CK2 target sites, we identified several serine residues in ZYG-1 that conform to the CK2 consensus motif (**Table S1**). Here, we focused on four serine sites, including S279 and adjacent serine residues (ZYG-1:4S) in the ZYG-1 L1 domain (**Fig 2A**), which is critical for centrosomal ZYG-1 loading and direct binding to SAS-6^17,57^, and well conserved in closely related nematodes (**Fig S2A**). Our functional analyses reveal that mutating four serine residues simultaneously produces more potent effects on ZYG-1 activity than the S279 single mutation, indicating that S279 and at least one additional serine in ZYG-1:4S are phosphorylated *in vivo*. Furthermore, genetic and mass spectrometry analyses suggest that CK2 has at least one target site in ZYG-1:4S *in vivo* (**Fig 5B**) and phosphorylates at least one serine between S279 and S280 *in vitro* (**Fig S2D**). These results support the notion that CK2 directly phosphorylates ZYG-1:4S, and phosphorylation of ZYG-1 at multiple sites, including CK2 target site(s), regulates ZYG-1 activity in centrosome assembly.

### The phosphorylation state of ZYG-1 regulates ZYG-1 stability

Our collective data support that the NP-ZYG-1 mutations mimic loss of CK2. The NP-ZYg-1 mutations restore centrosome duplication and embryonic viability to the hypomorphic *zyg-1(it25)* mutants, indicating hyperactive ZYG-1. Remarkably, the NP-ZYG-1 mutations lead to centrosome amplification, reminiscent of the phenotype caused by Plk4/ZYG-1 overexpression. Consistently, total ZYG-1 levels are significantly increased in the NP-ZYG-1 embryos, leading to elevated levels of ZYG-1 and downstream factors at centrosomes, providing a possible mechanism of the NP-ZYG-1 mutation to drive centrosome amplification.

Moreover, our results suggest that PM-ZYG-1 is stabilized when the 26S proteasome is inhibited, while NP-ZYG-1 is partially resistant to the 26S proteasome, supporting our model where site-specific phosphorylation of ZYG-1 regulates ZYG-1 stability through proteasomal degradation. Thus, the interplay between CK2 and a phosphatase, potentially PP2A^B55^ based on our analysis discussed below, appears to balance the phosphorylation state of ZYG-1, controlling ZYG-1 levels through proteolysis to limit the centrosome number.

### Regulation of CK2-dependent phosphorylation of ZYG-1

Since the kinase CK2 is constitutively active^62^, a refractory mechanism to counteract CK2 should be available for balancing the phosphorylation state of ZYG-1. One mechanism could be through dephosphorylation of ZYG-1 by protein phosphatase. In *C. elegans*, PP2A^SUR-6/B55^ positively regulates ZYG-1 levels^31^, similar to the proposed role of PP2A^Twins/B55^ in stabilizing Plk4 in *Drosophila*^32^. Serine residues in the ZYG-1:4S align with the PP2A^B55^ consensus motif surrounded by polybasic residues^59,60^ (**Fig 2A,S2A**), suggesting that PP2A^SUR-6/B55^ may stabilize ZYG-1 by counteracting CK2-dependent phosphorylation at ZYG-1:4S. Another possibility is through the substrate availability that restricts ZYG-1 phosphorylation. ZYG-1 undergoes stepwise interactions with centrosome proteins during centrosome biogenesis^17,57^. CK2 target sites in ZYG-1 may be masked or competed by other proteins, preventing CK2 targeting. Such a mechanism has been observed in human cells where Plk4 and CDK-1 compete for binding to STIL/Ana2/SAS-5, providing the temporal regulation of STIL phosphorylation^63^. Alternatively, efficient CK2 targeting may require conformational changes of ZYG-1. In human cells, Plk4 binding to STIL/Ana2/SAS-5 triggers Plk4 kinase activity^10^. Likewise, ZYG-1 binding to another centrosome factor could induce optimal conformation for CK2 targeting. As the ZYG-1:4S resides within the ZYG-1 binding domain to SAS-6^17^, ZYG-1 binding to SAS-6 or another centrosome factor may induce conformational changes of ZYG-1, promoting or blocking CK2 targeting.

### Mechanism of ZYG-1 Phosphorylation

The mechanism of ZYG-1 phosphorylation in *C. elegans* has been relatively unexplored compared to the extensively studied autophosphorylation of Plk4 in other organisms. Plk4 autophosphorylation in the L1 domain is crucial for its kinase activation and stability^18,19,23,24,64,65^, ensuring proper centrosome number. Our study identifies a cluster of serine residues (ZYG-1:4S) in the ZYG-1 L1 domain that regulates ZYG-1 stability, analogous to Plk4 autophosphorylation. Both Plk4 and ZYG-1 have a conserved SBM responsible for SCF^Slimb/βTrCP^-mediated proteolysis^18–20,23,24^. While Plk4 activates the SBM phosphodegron through autophosphorylation^19,23,24^, it remains unclear if ZYG-1 autophosphorylation provides a similar mechanism, although the ZYG-1 SBM plays a conserved role in regulating ZYG-1 stability^20^. The ZYG-1 SBM (aa334-339), located outside ZYG-1:4S (aa273-280), does not conform to the CK2 consensus motif, suggesting that CK2 is unlikely to phosphorylate the ZYG-1 SBM. As ZYG-1 is autophosphorylated *in vitro*^12^, it is tempting to speculate that autophosphorylation of the ZYG-1 SBM promotes SCF-mediated destruction through a conserved mechanism. Therefore, ZYG-1 phosphorylation within ZYG-1:4S may regulate its stability independently of the SCF^Slimb/βTrCP^-mediated pathway. In *C. elegans*, centrosomal ZYG-1 levels are regulated by the anaphase promoting complex/cyclosome (APC/C)^FZR-1^ E3 ubiquitin ligase and SCF^Slimb/βTrCP20^. In humans, Plk4 levels are also regulated by another E3 ubiquitin ligase, Mib1^66^, and TEC tyrosine kinase-dependent phosphorylation^67^, indicating multiple pathways involved in regulating Plk4/ZYG-1 stability. Thus, CK2-dependent phosphorylation of ZYG-1 likely provides an additional regulatory mechanism to fine-tune ZYG-1 activity in centrosome assembly.

Our study provides insights into the mechanism of ZYG-1 phosphorylation and the critical role of CK2 kinase activity in proper centrosome assembly. This study is the first to report that CK2 directly phosphorylates ZYG-1/Plk4 in any organism. Considering the divergence of *C. elegans* ZYG-1 from Plk4 orthologs in other organisms^68,69^, it would be intriguing to investigate whether CK2 also phosphorylates Plk4 orthologs in other species. In human cells, elevated CK2 activity has been observed in cancer cells and normal proliferating cells^46,47^, and dysregulation of CK2α activity has been linked to centrosome amplification. However, the specific targets involved in this process remain unknown^48^. Based on our findings, it is tempting to speculate that CK2 may influence centrosome assembly by phosphorylating Plk4, potentially through a similar regulatory mechanism observed in *C. elegans*. Understanding whether CK2 directly phosphorylates Plk4 and how aberrant CK2 activity relates to abnormal centrosome number in human cells remain important questions for future studies.

## Methods and Materials

### *C. elegans* Strains and Genetic Analysis

All strains used in this study were derived from the N2 wild-type strain^70,71^. Animals were maintained at 16 or 19°C on MYOB plates seeded with *Escherichia coli* OP50. For embryonic lethality assays, L4 animals were transferred to individual plates and allowed to produce progeny for 24-48 h. Progeny was then allowed to develop for 18-24 h before scoring the number of larva and unhatched (dead) embryos. RNAi feeding was performed as previously described^72^ using the L4440 empty vector as a negative control. Partial depletion of *rpt-4* was achieved by briefly feeding for 12-18 h before imaging. A complete list of *C. elegans* strains used in this study is provided in Table S3. Some strains were provided by the CGC, funded by the NIH Office of Research Infrastructure Programs (P40 OD010440).

### CRISPR/Cas9 Genome Editing and Microinjection

Microinjection was performed using the XenoWorks microinjector (Sutter Instruments, Novato, CA) with a continuous pulse setting at 400–800 hPa. For all genome editing, we used the *dpy-10(cn64)* co-CRISPR technique as previously described^73,74^. Animals were microinjected with Cas9 RNP complexes preloaded with equimolar crRNA (0.4-0.8 µg/µl each) and tracrRNA molecules (12 µg/µl) as previously described^75^. Single-stranded oligonucleotide repair templates were included at 25-100 ng/µl. Cas9 and custom-designed oligonucleotides (Tables S2, S3) were purchased from IDT (Coralville, IA). Following injection, F1 generation *dpy-10(cn64)* II/+ rollers were screened for each co-edit. All genotypes were verified in homozygous F2 generation animals through Sanger Sequencing (GeneWiz, South Plainfield, NJ).

### Expression and purification of recombinant proteins

pGEX-6P-1 plasmids for expressing full length ZYG-1 were provided by Genewiz and transformed into Rosetta 2(DE3) pLysS cells. Starter cultures were inoculated using 2L of LB media and grown at 37°C until the mixture reached an optical density of 0.4-0.6 at 600 nm. Protein expression was induced by adding 1 mM isopropyl ß-D-1-thiogalactopyranoside (IPTG) and incubating the cultures overnight at 16°C (250 rpm). The cells were harvested by centrifugation at 4,000 rcf, and the pellet was resuspended in Lysis Buffer [PBS with 250mM NaCl, 10mM EDTA, 0.1% Tween 20, 200 μg/ml Lysozyme, 0.1% PMSF]. The cells were lysed by sonication at 40% amplitude and a 10s/50s on/off pulse using a 505 Sonic Dismembrator equipped with a ¼” titanium probe (Fisher Scientific). The lysate was then clarified by centrifugation at 18,500 rpm for 1.5 hours using a Sorvall RC5C Plus centrifuge equipped with a SS34 rotor. The supernatant was then applied to a 10 mL packed GST resin (GE Healthcare, Waukesha, WI) equilibrated in Wash Buffer [PBS with 250 mM NaCl and 1 mM DTT]. The column was washed with 3x 20mL volumes of Wash Buffer, and the GST fusion proteins were eluted in 50 mM Tris [pH 8.1], 150 mM KCl, and 1mM DTT through a 5-50 mM glutathione gradient. The protein elutions were identified by SDS-PAGE and A280 absorbance, and the samples were dialyzed against Storage Buffer [20mM HEPES pH 7.5, 150mM KCl, 1mM DTT, and 40% Glycerol]. After three exchanges (4 hours each), the concentrated protein samples were snap-frozen in liquid nitrogen and stored at -80°C until use.

### *In vitro* Kinase Assays

Recombinant human Casein Kinase 2 (NEB, P6010S) and Protein Kinase A (NEB, P6000S) were purchased from New England Biolabs. Recombinant GST::ZYG-1^K41M^ (kinase-dead)^12,17^ fusion proteins were expressed in *Escherichia coli* and purified using glutathione resin. Protein concentration was determined using a nanodrop spectrophotometer. For kinase reactions using full-length ZYG-1, 1 μg of recombinant ZYG-1 proteins were incubated with 1 μg of recombinant CK2 in kinase buffer [500 mM Tris-Cl, 100 mM MgCl_2_, 1 mM EDTA and 20 mM DTT] for 30 minutes at room temperature. Following kinase reactions, proteins were analyzed by SDS/PAGE and autoradiography. ZYG-1 peptides were produced by Genscript (Piscataway, NJ). For ZYG-1 peptide analysis, 5-10 μg of ZYG-1 peptides were incubated at room temperature for 30 minutes with 1 μg of recombinant CK2 in kinase buffer [500 mM Tris-Cl, 100 mM MgCl_2_, 1 mM EDTA and 20 mM DTT]. Peptides were then analyzed through thin layer chromatography and autoradiography as previously described^76^. In brief, the phosphorylated peptides were loaded on silica plates and allowed to chromatograph for 4 h at room temperature. A mixture of n-butanol, acetic acid, pyridine, and water (3:1:2:4 ratio) was used as a solvent. For all *in vitro* kinase reactions, 200 μM of cold ATP and 4 μCi of ATP[γ-^32^P] were included in each reaction. To inhibit CK2 activity, 5 μM TBB (4,5,6,7-tetrabromobenzotriazole, Tocris, 2275) was added to reactions. For mass spectrometry, ZYG-1 peptides following *in vitro* kinase reaction without γ-^32^P were analyzed for the phosphorylation events by the Taplin Mass Spectrometry facility as described previously^31^.

### Immunofluorescence and Cytological Analysis

Confocal microscopy was performed as previously described^27^, using a Nikon Eclipse Ti-U microscope equipped with a Plan Apo 60× .4 NA lens, a Spinning Disk Confocal (CSU X1), and a Photometrics Evolve 512 camera. The following primary antibodies were used at 1:3000 dilutions: DM1a (Sigma, T9026), α-ZYG-1^56^, α-SAS-4^27^, α-Myc (ThermoFisher, PA1-981), α-V5 (MBL, M167-3), α-HA (ThermoFisher, 26183) and α-Ollas (ThermoFisher, MA5-16125). Alexa Fluor 488 and 568 secondary antibodies (ThermoFisher, A11001, A11004, A11006, A11034, A11036) were used at 1:3000 dilutions.

Image acquisition and quantification of fluorescence intensity were performed using MetaMorph software (Molecular Devices, Sunnyvale, CA, USA). Adobe Creative Cloud 2023 (Photoshop/Illustrator) was used for image processing. To quantify centrosomal signals, the average intensity within 8 or 9-pixel (1 pixel=0.151 µm) diameter region was recorded for a single focal plane within an area centered on the centrosome. The average intensity within a 25-pixel diameter region drawn outside the embryo was used for background subtraction.

### Immunoprecipitation (IP) and Western Blot

IP experiments were performed as described^56^. Embryos were extracted from adult worms using hypochlorite treatment (1:2:1 ratio of M9 buffer, 5.25% sodium hypochlorite, and 5 M NaOH), washed five times using M9 buffer, frozen in liquid nitrogen and stored at -80°C until use. Embryos were suspended in lysis buffer [50 mM HEPES, pH 7.4, 1 mM EDTA, 1 mM MgCl_2_, 200 mM KCl, and 10% glycerol (v/v)] supplemented with complete protease inhibitor cocktail (Roche, Basel, Switzerland) and MG132 (Tocris, Avonmouth, Bristol, UK). Embryos were then milled for 5 minutes (repeat x3) at 30 Hz using a Retsch MM 400 mixer-mill (Verder Scientific, Newtown, PA) and sonicated for 3 minutes in an ultrasonic water bath (Thermo Fisher, Waltham, MA).

Lysates were spun at 45,000 rpm for 45 minutes using a Sorvall RC M120EX ultracentrifuge (Thermo Fisher), then the supernatant was recovered to clean tubes. Protein concentrations were determined using a nanodrop spectrophotometer, and the equivalent amount of total protein lysates was used for IP. Embryonic protein lysates mixed with α-Myc magnetic beads (MBL, M047-11) were incubated by rotation for 1 hour at 4°C and washed (3x 5 minutes) with PBST (PBS + 0.1% Triton-X 100). IP with beads and input samples were resuspended in 2X Laemmli Sample Buffer (Sigma) and boiled for 5 minutes before fractionating on a 4–12% NuPAGE Bis-Tris gel (Invitrogen) and transferred to nitrocellulose membrane. The following antibodies were used at 1:3000-10,000 dilutions: DM1a (Sigma, T9026), α-V5 (Genscript, A01724), α-Myc (Genscript, A00704), α-SAS-6^31^, and IRDye secondary antibodies (LI-COR Biosciences). Blots were imaged using the Odyssey infrared scanner (LI-COR Biosciences) and analyzed using Image Studio software (LI-COR Biosciences).

## Supporting information

Medley et al 2023-Supplemental Files

## Statistical analysis

Statistics were generated using R statistical software and presented as average ± standard deviation (SD). Dotplots were generated using the R ‘beeswarm’ package. In the dotplots, boxes range from the first through third quartile of the data. Thick bar indicates the median. Solid grey line extends 1.5 times the inter-quartile range or to the minimum and maximum data point. All P-values were calculated using two-tailed t-tests: ^ns^*p*>0.05; \**p*<0.05; \*\**p*<0.01; \*\*\**p*<0.001.

