## Supplementary material for "Site-Specific Phosphorylation of ZYG-1 Regulates ZYG-1 Stability and Centrosome Number": Medley et al 2023-Supplemental Files

[4] Corresponding Author

**Table S1:** ***In silico*** **Prediction of CK2 Phosphorylation Sites on ZYG-1**

| Site | Sequence | Domain | GPS5.0 | KinasePhos3.0 |
|---|---|---|---|---|
| S2 | ******M**S**GGKSGSR | N-term | | 1.000 |
| S6 | **MSGGK**S**GSRLSAY | N-term | | 0.576 |
| S273 | RDGRRQR**S**REPVRSS | L1 | | 0.755 |
| **S279** | RSREPVR**S**SRDDRSR | L1 | **3.194** | **1.000** |
| S280 | SREPVRS**S**RDDRSRD | L1 | 2.302 | **1.000** |
| S285 | RSSRDDR**S**RDGRALI | L1 | | 0.607 |
| S294 | DGRALIR**S**SSQPAHS | L1 | | 0.770 |
| S295 | GRALIRS**S**SQPAHSG | L1 | | 0.500 |
| S296 | RALIRSS**S**QPAHSGR | L1 | | 0.541 |
| S301 | SSSQPAH**S**GRAPLSN | L1 | | 0.545 |
| S319 | HDRMPST**S**SRGFDSE | L1 | | 0.574 |
| S320 | DRMPSTS**S**RGFDSER | L1 | | 0.615 |
| S325 | TSSRGFD**S**ERGRERD | L1 | 1.325 | 0.986 |
| S335 | GRERDRD**S**GRGTVPP | L1 | | 0.638 |
| T339 | DRDSGRG**T**VPPSRED | L1 | | 0.630 |
| S343 | GRGTVPP**S**REDRNRS | L1 | | 0.922 |
| S573 | SPSSLMP**S**GSSQTSR | L2 | | 0.953 |
| T605 | APFLTKP**T**SSQRASS | L2 | | 0.707 |
| S606 | PFLTKPT**S**SQRASSA | L2 | | 0.801 |
| S607 | FLTKPTS**S**QRASSAN | L2 | | 0.760 |
| S611 | PTSSQRA**S**SANVQRR | L2 | | 0.989 |
| **S620** | ANVQRRV**S**TDENSSP | L2 | **5.516** | **1.000** |
| T621 | NVQRRVS**T**DENSSPS | L2 | 1.562 | **1.000** |
| S625 | RVSTDEN**S**SPSVAPS | L2 | | 0.958 |
| S626 | VSTDENS**S**PSVAPSK | L2 | | 0.831 |
| S628 | TDENSSP**S**VAPSKYK | L2 | | 0.648 |

Prediction scores for different in silico phosphorylation prediction programs examining the likelihood of site-specific, CK2-dependent phosphorylation of ZYG-1. The full-length *C. elegans* ZYG-1 protein (Wormbase version WS284) was submitted to each respective program. For GPS5.0 (Wang et al.[1]) predictions, the GPS5.0 webserver (http://gps.biocuckoo.cn/online.php) was used by selecting "CMCG > CK2" and using a low threshold. All serine/threonine residues scoring over the 0.184 cutoff for the CK2 kinase are presented under the GPS5.0 column as positive predictions. For KinasePhos3.0 (Ma et al[2]), the KinasePhos3.0 application was downloaded (https://awi.cuhk.edu.cn/KinasePhos/download.html) and run locally by selecting the "CSK21_Human" protein prediction. All sites given a prediction score by KinasePhos3.0 are shown in the table. The predicted serine or threonine residue is indicated by bolded font. Asterisks indicate areas before the first amino acid of ZYG-1 in the alignment.

**Table S2:** ***In silico*** **Prediction of CK2 Phosphorylation Sites on Human Plk4**

| Site | Sequence | Domain | GPS5.0 | KinasePhos3.0 |
|---|---|---|---|---|
| T34 | YRAESIH**T**GLEVAIK | Kinase | 0.210 | |
| S109 | KNRVKPF**S**ENEARHF | Kinase | 2.444 | 0.648 |
| S192 | RSAHGLE**S**DVWSLGC | Kinase | 1.147 | |
| S269 | HPFMSRN**S**STKSKDL | L1 | | 0.599 |
| S270 | PFMSRNS**S**TKSKDLG | L1 | | 0.638 |
| T271 | FMSRNSS**T**KSKDLGT | L1 | | 0.744 |
| S270 | PFMSRNS**S**TKSKDLG | L1 | 0.934 | |
| S273 | SRNSSTK**S**KDLGTVE | L1 | | 0.578 |
| S278 | TKSKDLG**T**VEDSIDS | L1 | | 0.845 |
| S298 | STAITAS**S**STSISGS | L1 | | 0.961 |
| S330 | TVFPKNK**S**STDFSSS | L1 | | 0.784 |
| T332 | FPKNKSS**T**DFSSSGD | L1 | | 0.892 |
| S335 | NKSSTDF**S**SSGDGNS | L1 | | 0.996 |
| S336 | KSSTDFS**S**SGDGNSF | L1 | **3.342** | **0.988** |
| S337 | SSTDFSS**S**GDGNSFY | L1 | | 0.650 |
| S352 | TQWGNQE**T**SNSGRGR | L1 | | 0.544 |
| S355 | GNQETSN**S**GRGRVIQ | L1 | | 0.612 |
| S378 | RYLRRAY**S**SDRSGTS | L1 | | 0.660 |
| S379 | YLRRAYS**S**DRSGTSN | L1 | 2.099 | 0.810 |
| S382 | RAYSSDR**S**GTSNSQS | L1 | | 0.997 |
| T384 | YSSDRSG**T**SNSQSQA | L1 | 0.237 | **1.000** |
| S385 | SSDRSGT**S**NSQSQAK | L1 | | **1.000** |
| S387 | DRSGTSN**S**QSQAKTY | L1 | | 0.996 |
| S389 | SGTSNSQ**S**QAKTYTM | L1 | | 0.941 |
| S408 | SAEMLSV**S**KRSGGGE | L1 | | 0.835 |
| S411 | MLSVSKR**S**GGGENEE | L1 | | 0.951 |
| S421 | GENEERY**S**PTDNNAN | L1 | 2.505 | 0.987 |
| T423 | NEERYSP**T**DNNANIF | L1 | | 0.544 |
| S438 | NFFKEKT**S**SSSGSFE | L1 | | 0.618 |
| S439 | FFKEKTS**S**SSGSFER | L1 | | 0.679 |
| S440 | FKEKTSS**S**SGSFERP | L1 | | 0.807 |
| S441 | KEKTSSS**S**GSFERPD | L1 | | 0.931 |
| S443 | KTSSSSG**S**FERPDNN | L1 | | 0.982 |
| S490 | LQINDPL**S**EQSKTRG | L1 | | 0.800 |
| S493 | NDPLSEQ**S**KTRGMEP | L1 | | 0.724 |
| S528 | DTKVKKN**S**DASDNAH | L1 | 1.075 | |
| S534 | KTKKAVV**S**ILDSEEV | L1 | | 0.668 |
| S568 | VFGSDPL**S**EQSKTRG | L1 | 3.140 | |
| S587 | PLADRPP**S**PTDNISR | CPB | | 0.719 |
| S641 | KEVLQIS**S**DGNTITI | CPB | 0.876 | |
| S665 | PLADRPP**S**PTDNISR | CPB | 2.072 | |
| S676 | GKSYTLK**S**ESEVNSL | CPB | | 0.893 |
| S678 | SYTLKSE**S**EVNSLKE | CPB | | 0.870 |
| S682 | KSESEVN**S**LKEEIKM | CPB | | 0.876 |
| S709 | LALESII**S**EEERKTR | CPB | | 0.981 |
| S718 | PLADRPP**S**PTDNISR | CPB | 1.097 | |
| S733 | GRKPGST**S**SPKALSP | CPB | | 0.908 |
| S734 | RKPGSTS**S**PKALSPP | CPB | | 0.675 |
| S739 | TSSPKAL**S**PPPSVDS | CPB | | 0.856 |
| S746 | SPPPSVD**S**NYPTRER | CPB | | 0.544 |
| S754 | GKSYTLK**S**ESEVNSL | CPB | 2.447 | |
| S760 | KSESEVN**S**LKEEIKM | CPB | 1.998 | |
| S787 | LALESII**S**EEERKTR | CPB | 2.372 | |
| S790 | LGLTTTA**S**GTDISSN | CPB | | 0.895 |
| T792 | LTTTASG**T**DISSNSL | CPB | | 0.849 |
| S795 | TASGTDI**S**SNSLKDC | CPB | | 0.615 |
| T851 | GVSSISY**T**SPNGQTT | L2 | | 0.538 |
| S852 | VSSISYT**S**PNGQTTR | L2 | | 0.615 |
| T858 | VSSISYT**S**PNGQTTR | L2 | | 0.677 |
| S868 | LGLTTTA**S**GTDISSN | L2 | 0.337 | |

Prediction scores for different in silico phosphorylation prediction programs examining the likelihood of site-specific, CK2-dependent phosphorylation of human Plk4. The full length human Plk4 protein (Uniprot O00444) was submitted to each respective program. For GPS5.0 (Wang et al.[1]) predictions, the GPS5.0 webserver (http://gps.biocuckoo.cn/online.php) was used by selecting “CMCG > CK2” and using a low threshold. All serine/threonine residues scoring over the 0.184 cutoff for the CK2 kinase are presented under the GPS5.0 column as positive predictions. For KinasePhos3.0 (Ma et al.[2]), the KinasePhos3.0 application was downloaded (https://awi.cuhk.edu.cn/KinasePhos/download.html) and run locally by selecting the “CSK21_Human” protein prediction. All sites given a prediction score by KinasePhos3.0 are shown in the table. The predicted serine or threonine residue is indicated by bolded font.

**Table S3. List of *C. elegans* Strains Used in this Study**

| Strain | Genotype | Origin |
|---|---|---|
| **N2** | *wild-type* | CGC |
| **OC14** | *zyg-1(it25[ZYG-1$^{P442L}$]) II* | O'Connell et al.[3] |
| **MTU21** | *zyg-1(mhs389it25[ZYG-1$^{4D:P442L}$]) II* | This Study |
| **MTU25** | *zyg-1(mhs399it25[ZYG-1$^{S279A:P442L}$]) II* | This Study |
| **MTU44** | *zyg-1(mhs404it25[ZYG-1$^{S279D:P442L}$]) II* | This Study |
| **MTU50** | *zyg-1(mhs409[ZYG-1$^{4D}$]) II* | This Study |
| **MTU86** | *zyg-1(mhs428it25[ZYG-1$^{4A:P442L}$]) II* | This Study |
| **MTU127** | *zyg-1(mhs454[ZYG-1$^{4A}$]) II* | This Study |
| **MTU137** | *kin-3(mhs464[KIN-3::V5]) I* | Yim et al.[4] |
| **MTU150** | *sas-6(mhs451[Ollas::SAS-6]) IV* | Medley et al.[5] |
| **MTU158** | *zyg-1(mhs454) II; sas-6(mhs451) IV* | This study |
| **MTU159** | *zyg-1(mhs409) II; sas-6(mhs451) IV* | This study |
| **MTU178** | *zyg-1(mhs456mhs482[2xMyc::ZYG-1$^{4D}$]) II* | This Study |
| **MTU209** | *zyg-1(mhs507mhs454[2xMyc::ZYG-1$^{4A}$]) II* | This Study |
| **MTU257** | *sas-5(mhs533[SAS-5::V5]) V* | Medley et al.[5] |
| **MTU271** | *zyg-1(mhs456[2xMyc::ZYG-1]) II* | Medley et al.[5] |
| **MTU299** | *zyg-1(mhs507mhs454) II; unc-119(ed3) III; bsIs15[pNP99:unc-119(+) tbb-1p::mCherry::tbb-2::tbb-2 3'UTR* | |
| **MTU337** | *spd-2(mhs580[SPD-2::2xHA]) I* | This Study |
| **MTU371** | *zyg-1(mhs454) II; sas-5(mhs533) V* | This study |
| **MTU373** | *zyg-1(mhs409) II; sas-5(mhs533) V* | This study |
| **MTU376** | *spd-2(mhs580) I; zyg-1(mhs454) II* | This study |
| **MTU377** | *spd-2(mhs580) I; zyg-1(mhs409) II* | This study |
| **MTU746** | *zyg-1(mhs456) II; unc-119(ed3) III; bsIs15[pNP99:unc-119(+) tbb-1p::mCherry::tbb-2::tbb-2 3'UTR* | This study |
| **MTU806** | *kin-3(mhs464) I; zyg-1(mhs456) II* | This study |
| **OC481** | *unc-119(ed3) III; bsIs15[pNP99:unc-119(+) tbb-1p::mCherry::tbb-2::tbb-2 3'UTR* | Medley et al.[6] |
| **SA250** | *tjIs54[pie-1p::GFP::tbb-2 + pie-1p::2xmCherry::tbg-1 + unc-119(+)]; tjIs57[pie-1p::mCherry::his-48 + unc119(+)]* | Toya et al.[7] |

**Table S4. List of crRNA for CRISPR/Cas9 Genome Editing**

| Gene | Target | Sequence (5'-3') | Origin |
|---|---|---|---|
| *dpy-10* | *dpy-10(cn64)* (co-CRISPR) | UUCUGCUGUCUUGAUUGACG | Arribere et al.[8] |
| *spd-2* | C-terminus | UCUAUUCGAAAAUCUUGUAU | This Study |
| *zyg-1* | N-terminus | UUGAUCAACUAUGAGAUGAG | This Study |
| *zyg-1* | ZYG-1-L1 | UGGACGACGACAGAGAUCGA | This Study |

**Table S5. List of ssODN Homologous Repair Templates for CRISPR/Cas9 Genome Editing**

| Gene | Variation | Sequence (5'-3') |
|---|---|---|
| *dpy-10* (Arribere et al.[8]) | *dpy-10(cn64)* | CACTTGAACTTCAATACGGCAAGATGAGAATGACTGGA AACCGTACCGCATGCGGTGCCTATGGTAGCGGAGCTTC ACATGGCTTCAGACCAACAGCCTAT |
| *spd-2* | C-terminal 2xHA Tag | AATCAGACATTTGTCAACGACGTTACAATTGTTCCGAAT ACAAGATTTTCGAATAGAAAGGGAGGTTCCGGTGGATC TGGTGGATCCTACCCATACGATGTTCCAGATTACGCTTA TCCATATGATGTTCCAGATTATGCTTAAAATCTTAACCTA ACTTTCCAAATATTCTCTG |
| *zyg-1* | N-terminal 2xMyc Tag | ATACGCTGTGCGAACGTATGAGAAAACTACATCAAGGT GGAAGTGGTGGCTCGGGTGGCTCTTACCCTTATGACGT ACCAGATTACGCGTATCCATACGATGTCCCTGATTACGC ATAATAGTAATAACTTAGTATACGATGAGTTTTGCTCTTA CTTCATGTGCTCCTACATTTTACCACATA |
| *zyg-1* | S279A | AGAACACTCGCGGGATGGACGACGACAGAGATCGCGA GAACCAGTAAGAGCCTCAAGAGATGATCGATCTCGAGA TGGCAGAGCTCT |
| *zyg-1* | S279D | AGAACACTCGCGGGATGGACGACGACAGAGATCGCGA GAACCAGTAAGAGACTCAAGAGATGATCGATCTCGAGA TGGCAGAGCTCT |
| *zyg-1* | 4A | CTTCTCGAGAGAACACTCGCGGGATGGACGACGACAG AGAGCCAGAGAACCAGTACGTGCCGCCAGAGATGATC GAGCCCGAGATGGCAGAGCTCTGATAAGGTCTTCGAGT CAACCTGC |
| *zyg-1* | 4D | CTTCTCGAGAGAACACTCGCGGGATGGACGACGACAG AGAGACAGAGAACCAGTACGTGACGACAGAGATGATCG AGATCGAGATGGCAGAGCTCTGATAAGGTCTTCGAGTC AACCTGC |

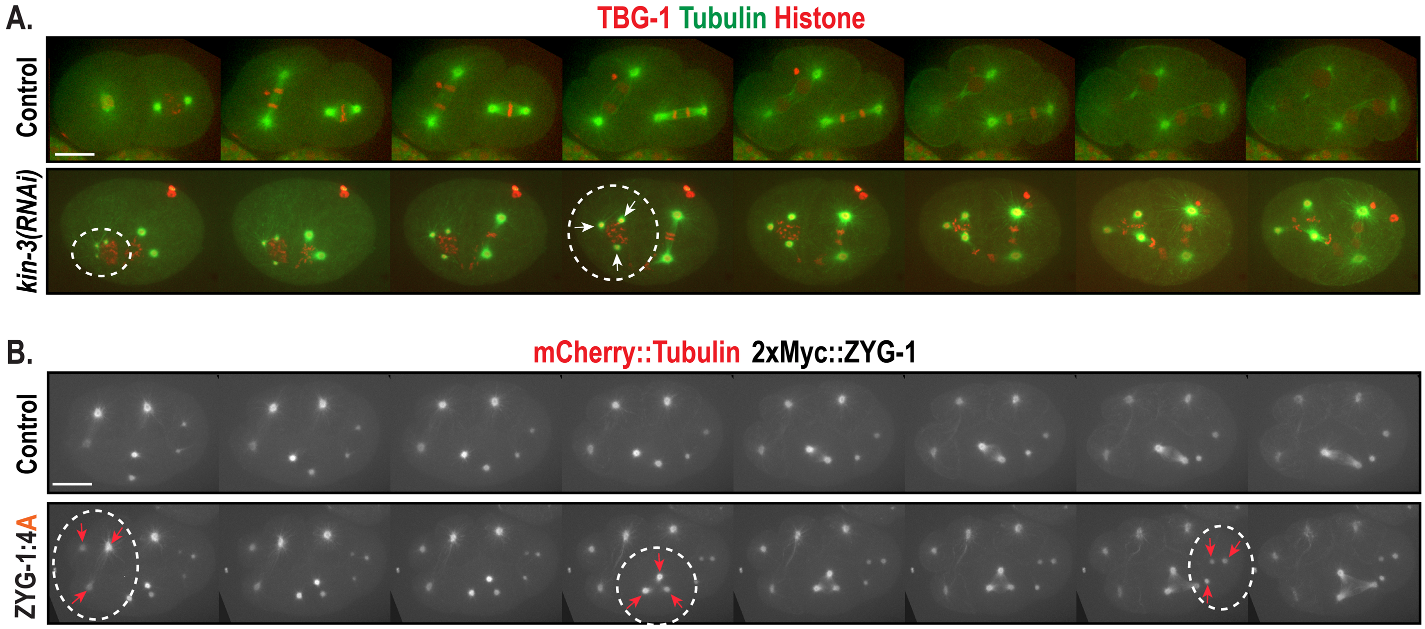


**Fig S1. Depleting CK2/KIN-3 or NP-ZYG-1:4A mutation leads to centrosome amplification.** Selected still images from 90s-interval timelapse movies of **(A)** embryos expressing GFP::β-tubulin, mCherry::TBG-1(γ-tubulin) and mCherry::histone, **(B)** 2xMyc::ZYG-1 embryos expressing mCherry::Tubulin. Arrows highlight tripolar spindles formed (circled).

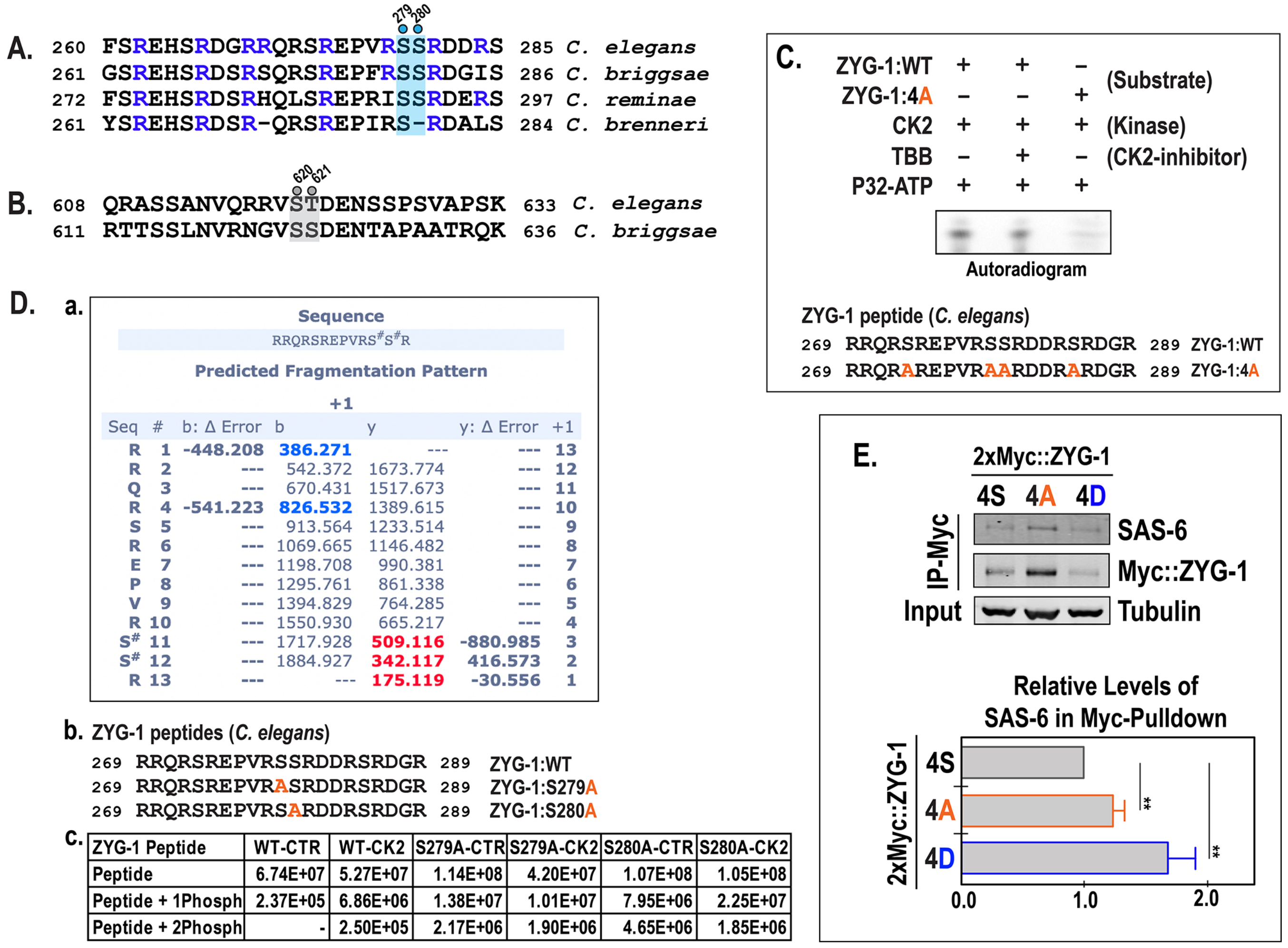


| ZYG-1 Peptide | WT-CTR | WT-CK2 | S279A-CTR | S279A-CK2 | S280A-CTR | S280A-CK2 |
|---|---|---|---|---|---|---|
| Peptide | 6.74E+07 | 5.27E+07 | 1.14E+08 | 4.20E+07 | 1.07E+08 | 1.05E+08 |
| Peptide + 1Phosph | 2.37E+05 | 6.86E+06 | 1.38E+07 | 1.01E+07 | 7.95E+06 | 2.25E+07 |
| Peptide + 2Phosph | - | 2.50E+05 | 2.17E+06 | 1.90E+06 | 4.65E+06 | 1.85E+06 |

**Fig S2. CK2 likely phosphorylates the ZYG-1:4S *in vitro* (A)** The ZYG-1:4S region is well conserved in the *Caenorhabditis* species. S279 and S280 are marked (blue dots and highlighted). This region coincides with the domain that binds to SAS-6 (Lettman et al.[9]). The basic R (blue) residues essential for SAS-6 binding are conserved. Note that serine residues in the ZYG-1:4S conform to both CK2 and PP2A$^{B55}$ consensus motifs. **(B)** S620 and T621 (gray highlighted) are conserved between *C. elegans* and *C. briggsae*, but absent in *C. reminae* and *C. brenneri*. **(C)** Autoradiogram of thin-layer chromatography suggests that CK2 phosphorylates ZYG-1:WT peptide *in vitro,* not ZYG-1:4A. The CK2 inhibitor TBB reduces signal. **(D)** Mass spectrometry (MS) analysis of the ZYG-1 peptides following *in vitro* kinase (CK2) reactions: **a.** The ZYG-1:WT peptide is phosphorylated by CK2 *in vitro*, with the most probable sites to be S279[S11] and S280[S12]. However, MS data could not confidently assign the phosphorylation site between two serines. **b.** The ZYG-1 peptides used for MS analysis. **c**. The numbers are the relative peak height for control and CK2-treated ZYG-1 peptides. Assuming that the ionization efficiency is similar between the peptides with and without phosphate, the ratio of phosphorylation level was compared. In the WT-CTR, there is much less than one percent of the peptide with one phosphate, while in WT-CK2, it is roughly 11% as the peptide with one phosphate, suggesting that at least one site is phosphorylated. To evaluate whether CK2 phosphorylates with a preference one of the two serines, the mutant peptides (S279A and S280A) were analyzed, finding no difference in signal between the two mutant peptides. If the CK2 favors one of the two, the signal would be much stronger in one of the mutants. **(E)** Quantitative immunoblot of 2xMyc::ZYG-1 immunoprecipitation (IP). The signal of SAS-6 in Myc-IP were normalized to that of Myc::ZYG-1 IP (mean±s.d., n=3) . Both Myc-pulldowns of 4A and 4D show the ratio of SAS-6/2xMyc::ZYG-1 higher than controls. Tubulin was used as a loading control. ~5% of total embryonic lysates were loaded in input lanes. ***p*<0.01 (two-tailed unpaired *t*-tests).

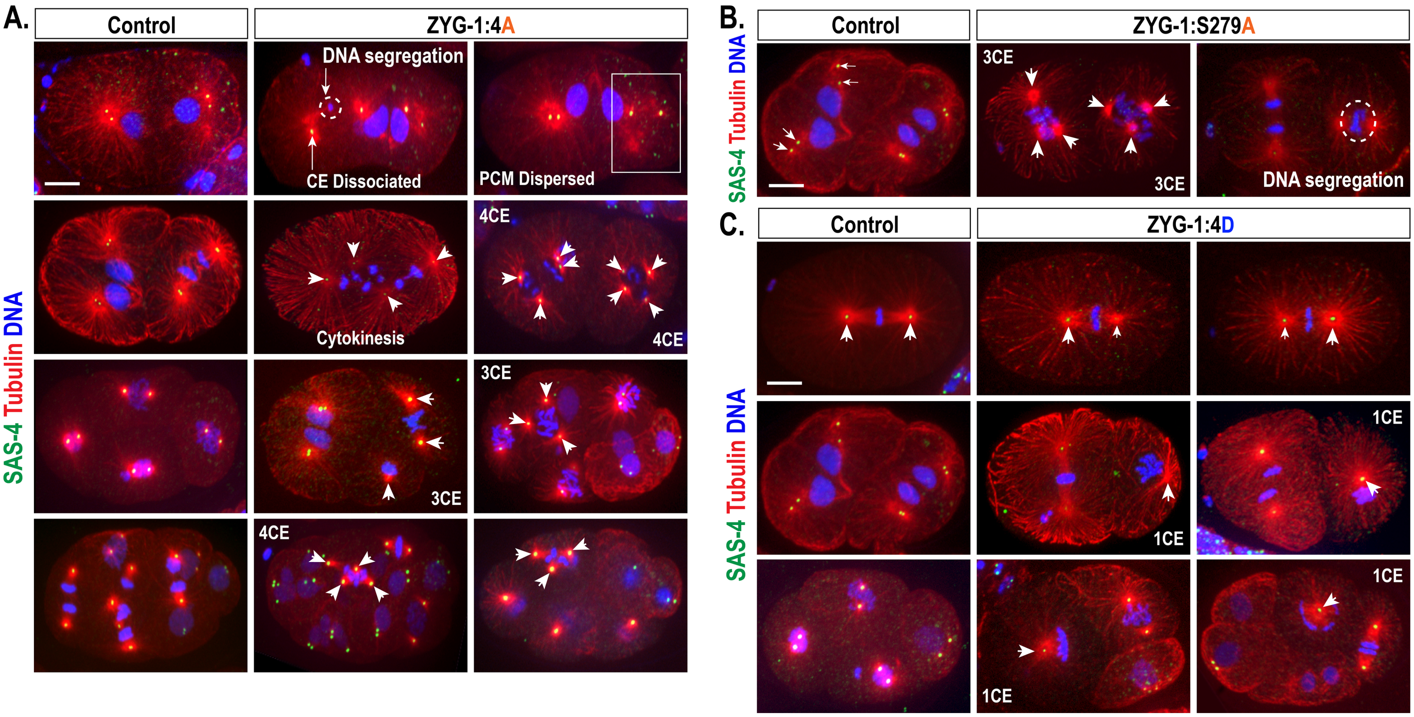


**Fig S3. ZYG-1 phospho-mutant embryos exhibit abnormal cell division phenotypes. (A)** Embryos stained for microtubules, SAS-4 and DNA illustrate various cell division phenotypes in ZYG-1:4A mutants, including DNA missegregation (circled), centrosome (CE) dissociation during late telophase, dispersed PCM, and cytokinesis failure. Supernumerary centrosomes are observed in later mitotic divisions (arrow heads). Tripolar mitotic spindles (3CE) strongly indicate centrosome amplification. **(B)** Examples of phenotypes observed in ZYG-1:S279A mutant embryos include extra centrosomes and multipolar spindle formation (arrowheads) and chromosome missegregation (circled). **(C)** Phenotypes observed in ZYG-1:4D mutant embryos. Asymmetric centrosomes, which reflect a partial block to centrosome duplication, are observed in ZYG-1:4D mutants (centrosome size reflected by arrowhead sizes). We also observe monopolar spindle formation (1CE) in ZYG-1:4D mutants, resulting from centrosome duplication failure (single arrowheads mark monopolar mitotic spindles). Bar, 10μm.

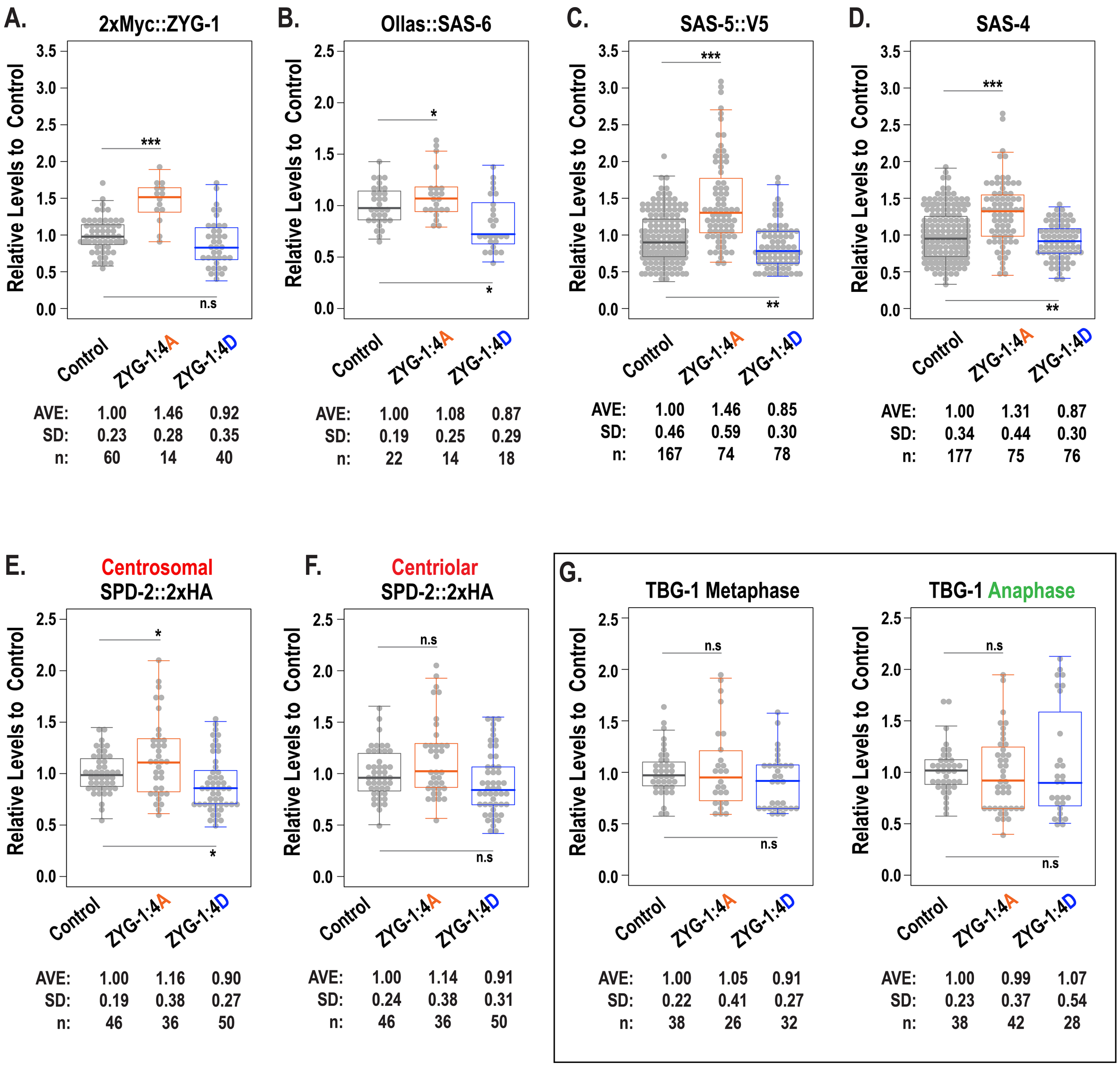


**Fig S4. Quantification of centrosomal factors at the first mitotic metaphase in ZYG-1 phospho-mutants. (A)** 2xMyc::ZYG-1, **(B)** Ollas::SAS-6, **(C)** SAS-5::V5, **(D)** SAS-4, **(E,F)** SPD-2::2xHA, and **(G)** centrosomal levels of TBG-1 remain unaffected in ZYG-1 phospho-mutants at the first mitotic metaphase and anaphase. ns $p>0.05$, *$p<0.05$, **$p<0.01$, ***$p<0.001$ (two-tailed unpaired *t*-tests). Boxes ranges from the first through third quartile of the data. Thick bars indicate the median. Lines extend to the minimum and maximum data point excluding outliers that were defined as beyond 1.5 times the interquartile range.
